# Methylation of Dual Specificity Phosphatase 4 Controls Cell Differentiation

**DOI:** 10.1101/2020.12.16.422727

**Authors:** Hairui Su, Ming Jiang, Chamara Senevirathne, Srinivas Aluri, Tuo Zhang, Han Guo, Juliana Xavier-Ferrucio, Shuiling Jin, Ngoc-Tung Tran, Szu-Mam Liu, Chiao-Wang Sun, Yongxia Zhu, Qing Zhao, Yuling Chen, LouAnn Cable, Yudao Shen, Jing Liu, Cheng-Kui Qu, Xiaosi Han, Christopher A. Klug, Ravi Bhatia, Yabing Chen, Stephen D. Nimer, Y. George Zheng, Camelia Iancu-Rubin, Jian Jin, Haiteng Deng, Diane S. Krause, Jenny Xiang, Amit Verma, Minkui Luo, Xinyang Zhao

**Affiliations:** Department of Biochemistry and Molecular Genetics, School of Medicine, the University of Alabama at Birmingham, Birmingham, Alabama, USA; Chemical Biology Program; Memorial Sloan Kettering Cancer Center, New York, New York, USA; Program of Pharmacology, Weill Cornell Medical College of Cornell University, New York, New York, USA; Department of Oncology, Albert Einstein Medical College, Montefiore Medical Center, Bronx, New York, USA; Genomics and Epigenomics Core Facility, Weill Cornell Medical College of Cornell University, New York, New York, USA; Tri-Institutional PhD Program in Chemical Biology, Memorial Sloan Kettering Cancer Center, New York, New York, USA; Department of Laboratory Medicine, Yale Stem Cell Center, Yale School of Medicine, New Haven, Connecticut, USA; Department of Medicine, School of Medicine, the University of Alabama at Birmingham, Birmingham, Alabama, USA; School of Life Sciences, Tsinghua University, Beijing, China; Array BioPharma, Boulder, Colorado, USA; Mount Sinai Center for Therapeutics Discovery, Departments of Pharmacological Sciences and Oncological Sciences, Tisch Cancer Institute, Icahn School of Medicine at Mount Sinai, New York, NY, USA; Department of Pediatrics, Emory School of Medicine; Aflac Cancer and Blood Disorders Center, Winship Cancer Institute, Emory University, Atlanta, Georgia, USA; Department of Neurology, School of Medicine, the University of Alabama at Birmingham, Birmingham, Alabama, USA; Department of Microbiology, School of Medicine, the University of Alabama at Birmingham, Birmingham, Alabama, USA; Division of Hematology and Oncology, School of Medicine, School of Medicine, the University of Alabama at Birmingham, Birmingham, Alabama, USA; Department of Pathology; School of Medicine, the University of Alabama at Birmingham, Birmingham, Alabama, USA; Veterans Affairs Birmingham Medical Center, Research Department, Birmingham, Alabama, USA; Sylvester Comprehensive Cancer Center, University of Miami, Miami, Florida, USA; Department of Pharmaceutical and Biomedical Sciences, College of Pharmacy, University of Georgia, Athens, Georgia, USA; Department of Medicine, Hematology and Oncology Division, Icahn School of Medicine at Mount Sinai, New York, NY, USA

**Author notes:** These authors made equal contribution (Hairui Su, Ming Jiang, Chamara Senevirathne). The current affiliation: Division of Hematology/Oncology, Boston Children’s Hospital, Harvard Medical School, Boston (Juliana Xavier-Ferrucio). Corresponding Authors: Xinyang Zhao,; Minkui Luo,; Amit Verma.

## Abstract

A collection of signaling and epigenetic events needs to be orchestrated for normal development of hematopoietic lineages. While mitogen-activated protein (MAP) kinases (MAPKs) and multiple epigenetic modulators have been implicated in the megakaryocytic (Mk) cell differentiation, the underlying molecular mechanisms of signaling-epigenetic crosstalk remain unclear. MAPKs are in general inactivated by dual specificity phosphatases (DUSPs), whose activities are tightly regulated by various posttranslational modifications. Using knockdown screening and single-cell transcriptional analysis, we determined that DUSP4 is the phosphatase that inactivates p38 MAPK in hematopoietic cells and serves as a key regulator to promote Mk differentiation. With the nextgeneration Bioorthogonal Profiling of Protein Methylation technology for live cells, we identified DUSP4 as a PRMT1 substrate. Mechanistically, PRMT1-mediated Arg351 methylation of DUSP4 triggers its ubiquitinylation by HUWE1 (an E3 ligase) and then degradation, which results in p38 MAPK activation and inhibition of Mk differentiation *in vitro* and *in vivo*. Interestingly, the mechanistic axis of the DUSP4 degradation and p38 activation is also associated with a transcriptional signature of immune activation and thus argues immunological roles of Mk cells. Collectively, these results demonstrate a critical role of PRMT1-mediated posttranslational modification of DUSP4 in regulation of Mk differentiation and maturation. In the context of thrombocytopenia observed in myelodysplastic syndromes (MDS), we demonstrated that high levels of p38 MAPK and PRMT1 are associated with low platelet counts and adverse prognosis, while pharmacological inhibition of p38 MAPK or PRMT1 stimulates megakaryopoiesis in MDS samples. These findings provide novel mechanistic insights into the role of the PRMT1-DUSP4-p38 axis on Mk differentiation and present a targeting strategy for treatment of thrombocytopenia associated with myeloid malignancies such as MDS.

## Introduction

Precisely orchestrated cellular differentiation is essential for normal development of a multicellular organism (1). Protein posttranslational modifications such as phosphorylation and methylation involved in the combination of signal transduction and epigenetic regulation are essential for cellular differentiation (2). Conversely, erasing posttranslational modification marks attenuates or even reverses many signaling and epigenetic events. Mitogen-activated protein (MAP) kinases (MAPKs) are activated through phosphorylation of the TxY motif on their catalytic sites. MAPK signaling is in general inactivated by a collection of dual specificity phosphatases (DUSPs), which function by dephosphorylating both threonine and tyrosine residues in the TxY motif, leading to inactivation of MAPK signaling (3-6). Furthermore, the activity of DUSPs is tightly regulated by posttranslational modifications such as phosphorylation and acetylation (5, 7).

Hematopoietic stem cells (HSCs) give rise to multiple types of lineage-specific progenitor cells with distinct gene expression patterns. A differentiation hierarchy of HSCs to erythroid (Er) and megakaryocytic (Mk) cells has been described through short-term HSCs, multipotent progenitors (MPPs), common myeloid progenitors (CMPs), and megakaryocyte-erythroid progenitors (MEPs) according to characteristic surface markers and transcriptional signatures (reviewed in (1)). Nevertheless, recent single-cell RNA sequencing (scRNA-seq) analysis of human CD34^+^ cells reveals a heterogeneous nature within surface marker-defined progenitors for gradually-evolved transcriptional expression topology during differentiation of hematopoietic stem/progenitor cells (HSPCs)(8, 9). Thus, extended data of essential signaling and epigenetic events, i.e. the so-called “hidden variables” (8), in addition to single-cell transcriptome snapshots, is necessary to define potential borders between lineage-specific progenitor cells and classify differentiated hematopoietic cells into functional subtypes. Activation of distinct MAPK cascades has been implicated in differentiation regulation of Er and Mk cells. A number of studies have shown that ERK promotes Mk differentiation and p38 MAPK activation promotes Er differentiation after stimulation by various cytokines (10-18). However, the precise molecular mechanisms that regulate the activation of MAPKs and phosphatases during Mk differentiation into mature Mk cells have not been fully elucidated.

PRMT1, one of the nine members of the human protein arginine methyltransferase (PRMT) family, has been associated with diverse biological functions such as transcriptional regulation, RNA processing, signal transduction, and DNA repair (19). In hematopoietic differentiation, PRMT1 is expressed at low levels in HSCs and is highly expressed in MEPs (20). Constitutive expression of PRMT1 reduces the generation of CD41^+^CD42^+^ Mk cells, presumably through arginine methylation of essential substrate(s) during Mk differentiation. One challenge to identifying PRMT1 substrates *in vivo* has been the lack of specific antibodies and the limitation of mass spectroscopy to detect methylarginine-containing peptides. To readily probe the activity of PRMTs, we developed the next-generation Bioorthogonal Profiling of Protein Methylation (BPPM) technology for live cells (21-24). Here, the cofactor-binding pockets of individual PRMTs were engineered to accommodate terminal-alkyne-containing bulky SAM analogues. The bulky SAM analogues were synthesized in live cells by an engineered methionine adenosyltransferase (MAT) with cell-permeable methionine analogues and endogenous adenosine triphosphate as substrates. The engineered PRMTs can then label substrates with a terminal-alkyne moiety from the cofactor analogues inside live cells. Using the live-cell BPPM technology, we identified that dual specificity phosphatase 4 (DUSP4 or MKP2, MAPK phosphatase 2) is a bona fide substrate of PRMT1 in broad cellular contexts.

In the context of Mk lineage determination, we show that PRMT1 triggers Arg351 methylation-dependent ubiquitinylation by the E3 ligase HUWE1 and thus DUSP4’s degradation, which result in activation of p38 MAPK signaling due to the inability of DUSP4 to dephosphorylate p38 MAPK. scRNA-seq analysis further confirmed the positive correlation between high levels of PRMT1 and the expression of signature genes of p38 activation including genes associated with pro-inflammatory response. We further show that pharmacological inhibition of p38 MAPK or PRMT1 promotes Mk differentiation both *in vivo* and *ex vivo*. In summary, we have identified a novel molecular crosstalk between signaling and epigenetic modulators that is critical for differentiation of Mk cells. These findings suggest potential pharmacological strategies for treatment of a commonly observed Mk cell differentiation defect in patients with myelodysplastic syndromes.

## Results

### Identification of DUSP4 as a key regulator for Mk differentiation

Distinct MAPK cascades regulate differentiation and homeostasis during hematopoiesis (25, 26). While MAPK cascades are modulated by DUSPs in general (27), the regulatory DUSP(s) specific for MAPK signaling during megakaryopoiesis remain to be elucidated. To address this question, we developed an shRNA-based screen to identify essential DUSP(s) regulating Mk differentiation (**Fig. 1a**). Here, we targeted 10 DUSPs known to modulate the signaling of MAPKs (5) and examined their specific roles in Mk differentiation. Given potential differences between factors regulating Mk cell differentiation of bone marrow (BM) cells and fetal liver cells (28), we tested both adult BM- and cord blood (CB)-derived human CD34^+^ HSPCs for their ability to differentiate into the Mk lineage when the individual DUSPs were downregulated using lentivirus-expressed shRNAs (**Fig. 1b**). In the presence of thrombopoietin (TPO), CD34^+^ BM and CB cells differentiated into mature Mk cells as characterized by CD41a^+^ and CD42b^+^ expression (**Fig. 1c** and **1d**). Upon knocking down specific DUSPs, only perturbation of DUSP4 significantly inhibited differentiation to CD41a^+^CD42b^+^ megakaryocytes (~40% reduction of the CD41a^+^CD42b^+^ population, **Fig. 1b** and **1c**). This requirement for DUSP4 during Mk differentiation was independently validated using BM and CB CD34^+^ cells from different donors (**Fig. 1d**), human G-CSF-mobilized peripheral blood (PB) CD34^+^ cells, and two additional DUSP4 shRNAs (**Fig. 1e-g**). While it is challenging to target Mk differentiation in a temporal manner (early progenitor cells, Mk progenitor cells, or matured Mk cells with different degrees of polyploidies), our results collectively argue the general requirement of DUSP4 for optimal Mk differentiation.

**Fig. 1.**
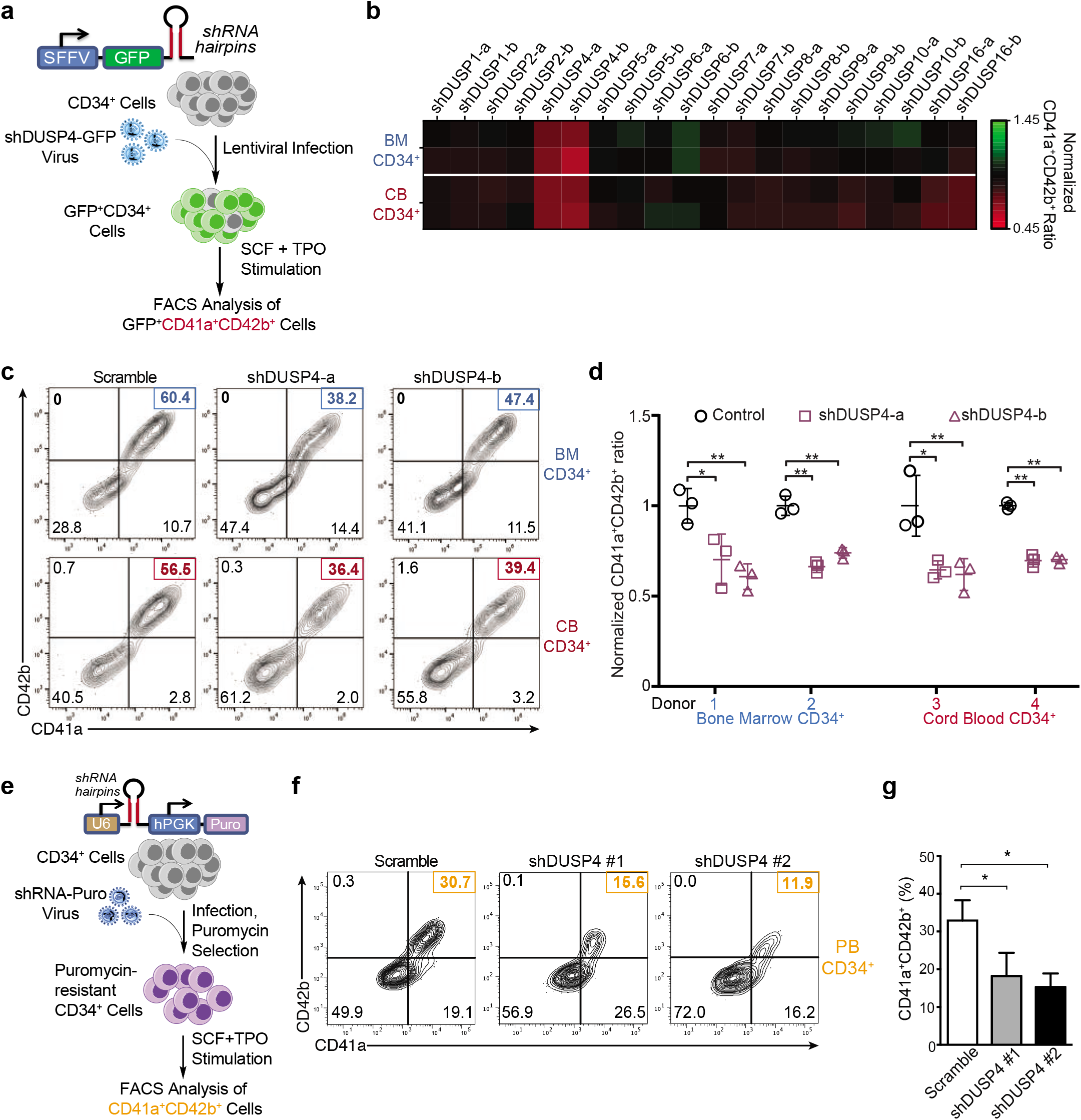
Identification of DUSP4 for optimal Mk differentiation. **a.** Schematic of shRNA-based screening assay to identify essential DUSP(s) for Mk-induced Mk differentiation. Human CD34^+^ cells isolated from bone marrow (BM) or cord blood (CB) were infected with lentiviruses expressing a control shRNA hairpin or shRNAs against DUSPs. Infected cells coexpressing GFP were sorted and cultured in the presence of TPO and SCF to stimulate Mk differentiation. Percentage of CD41a^+^CD42b^+^ cells was determined by FACS after 8-day culture. **b.** Heatmap of the percentages of CD41a^+^CD42b^+^ cells upon DUSP knockdown. Fold changes were normalized to the percentage of double positive cells with the group treated with control shRNA. **c.** Representative flow chart of FACS analysis of Mk differentiation using BM cells (top panel) and CB cells (bottom panel) cultured in TPO-containing medium. **d.** Summary of FACS analysis. Statistics are based on the data of 3 replicates (n = 3) with the BM or CB cells from two donors. **e.** Schematic description of Mk differentiation of human peripheral blood-derived CD34^+^ cells with DUSP4 knockdown. G-CSF mobilized peripheral blood CD34^+^ cells were infected with shRNA-encoding lentiviruses. Puromycin-selected cells were cultured in the presence of SCF and TPO for Mk differentiation. Percentage of CD41a^+^CD42b^+^ cells was determined by FACS after 7 days. **f.** and **g.** Representative and complete FACS analysis of CD41a and CD42b markers for Mk differentiation with peripheral blood CD34^+^ cells upon DUSP4 knockdown. Representative plots **f.** and statistics **g.** are shown (n = 3). Data were shown as mean ± SD, two-tailed paired t-test, *P ≤ 0.05; **P ≤ 0.01.

### DUSP4 drives the differentiation toward Mk progenitors

Megakaryocytes and erythrocytes can originate from the same progenitor subset like MEPs (29) or from even earlier stem/progenitor cells (30-32). We knocked down DUSP4 in human CD34^+^ cells grown in media containing both TPO and erythropoietin (EPO), which support differentiation to Mk or Er cells, respectively (**Fig. 2a**). The results showed that DUSP4 knockdown decreased differentiation efficiency to CD41a^+^ Mk cells and increased commitment to CD71^+^ Er cells (**Fig. 2b**). On average, we observed up to a two-fold reduction of CD41a^+^ megakaryocytes and a 30~50% increase in CD71^+^ erythrocytes (**Fig. 2b**). The differentiation bias observed with DUSP4 knockdown closely resembled what was observed when PRMT1 was overexpressed in CD34^+^ cells (20, 33).

**Fig. 2.**
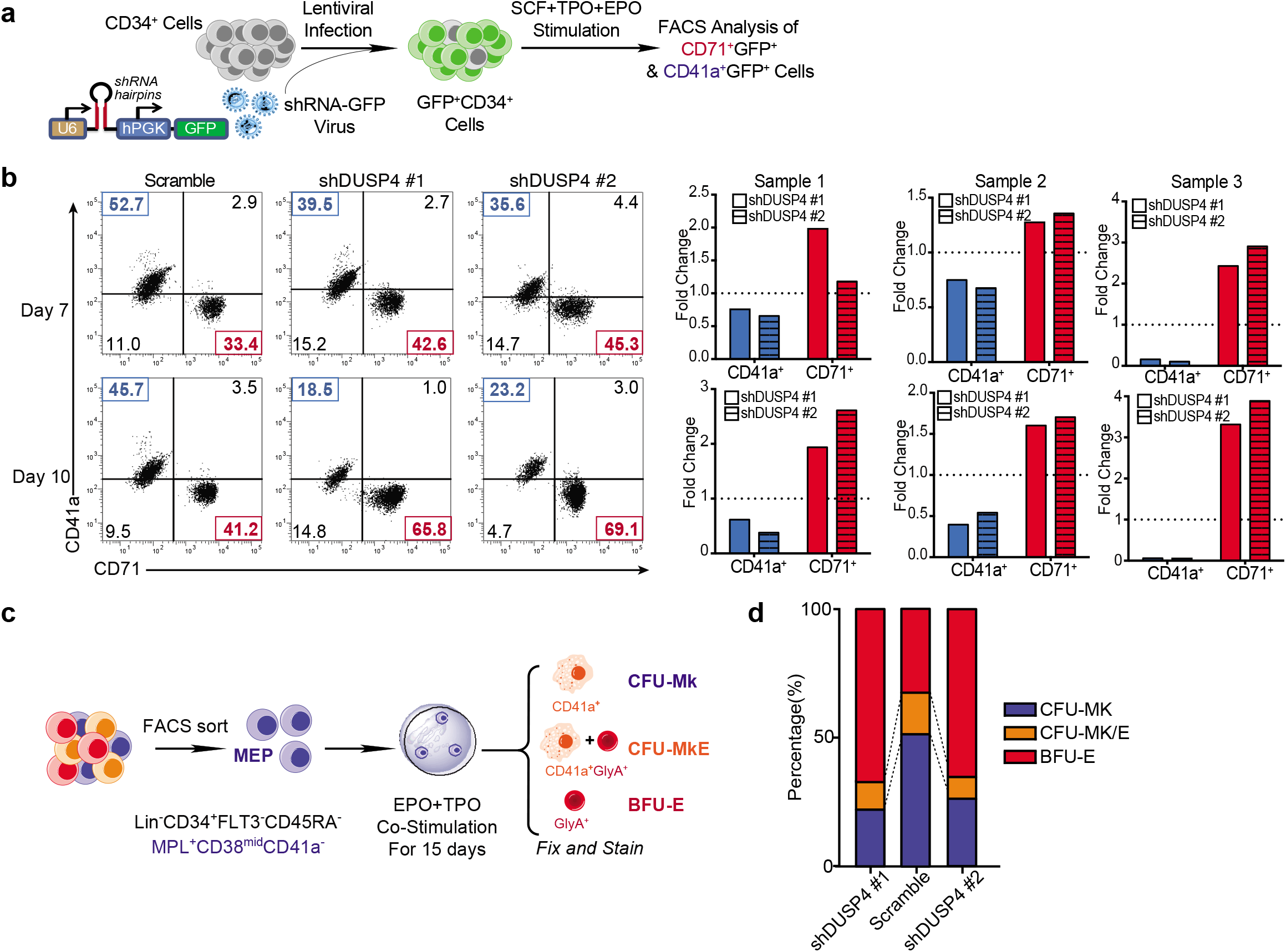
DUSP4-regulated differentiation choices between Mk cells and Er cells. **a.** Schematic of differentiation experiments using cord blood CD34^+^ cells. Cells were infected with shRNA lentiviruses and selected with GFP by flow cytometry. SCF/TPO/EPO were then added to induce differentiation to Mk and Er lineages. Percentages of CD41a^+^ and CD71^+^ cells were determined by FACS after 7 and 10 days. **b.** FACS analysis of CD41a and CD71 on the cultured cells. Representative plots (top panel) and normalized statistics of three samples (bottom panel) are shown (n=3). Fold changes were normalized with scramble controls. **c.** Schematic description of colony-forming unit (CFU) assays using different types of progenitor cells. Human CD34^+^ cells were infected with shRNA lentiviruses and then sorted by GFP together with respective markers for MEP, MkP, and ErP. The resulting cells were applied for colony-forming unit assays in the presence of EPO and TPO for Mk and Er differentiation. Colonies were characterized and counted after 14 days. **d.** Percentages of CFU-Mk, BFU-E, and CFU-MkE in each sorted population (n=2).

To explore the potential role of DUSP4 in regulating differentiation of MEP cells into Mk or Er cells (29, 34-36), we then isolated potential human MEPs, according to their respective cell surface marker profile: Lin^-^ CD34^+^FLT3^-^CD45RA^-^MPL^+^CD38^mid^CD41a^-^ for MEPs (**Fig. 2c**) (36) and performed colony-forming unit (CFU) assays, following transduction with the lentivirus-expressed DUSP4 shRNAs. Using MEPs, DUSP4 knockdown significantly increased BFU-E (burst-forming unit of erythrocytes by 2~3 fold) and decreased CFU-Mk (CFU of megakaryocytes by ~2-fold) (**Fig. 2d**). These results demonstrate that DUSP4 drives differentiation from biopotent MEP to Mk progenitor cells in colony formation assays.

### Specific deactivation of p38 MAPK by DUSP4 during Mk terminal differentiation

The human M7 leukemia cell line MEG-01, which represents immature Mk progenitors, can be stimulated to undergo differentiation and polyploidization with PMA (phorbol 12-myristate 13-acetate) treatment. We then used this system to examine the impact of DUSP downregulation on PMA-stimulated Mk terminal differentiation in cells that were transduced with the anti-DUSP shRNA library (**Extended Data Fig. 1a**). Consistent with observations using human BM, CB, and PB CD34^+^ cells (**Fig. 1**), only DUSP4 knockdown inhibited terminal Mk differentiation (~ 40% reduction of efficiency, **Extended Data Fig. 1b and 1c**). During the PMA-stimulated Mk differentiation of MEG-01 cells, *DUSP4* mRNA increased rapidly and reached a plateau by day 2 (**Fig. 3a**). In contrast, DUSP4 protein levels gradually increased over the course of differentiation, indicating a positive correlation between DUSP4 protein levels and Mk differentiation (**Fig. 3b**). We then examined whether DUSP4 overexpression would promote PMA-induced Mk differentiation of MEG-01 cells. Consistent with the necessity of DUSP4 for optimal Mk differentiation of human CD34^+^ HSPCs, DUSP4 overexpression directly increased the frequency of the CD41a^+^ population and led to a higher degree of polyploidy (**Extended Data Fig. 1d**). These data further confirm the essential role of DUSP4 in Mk maturation.

**Fig. 3.**
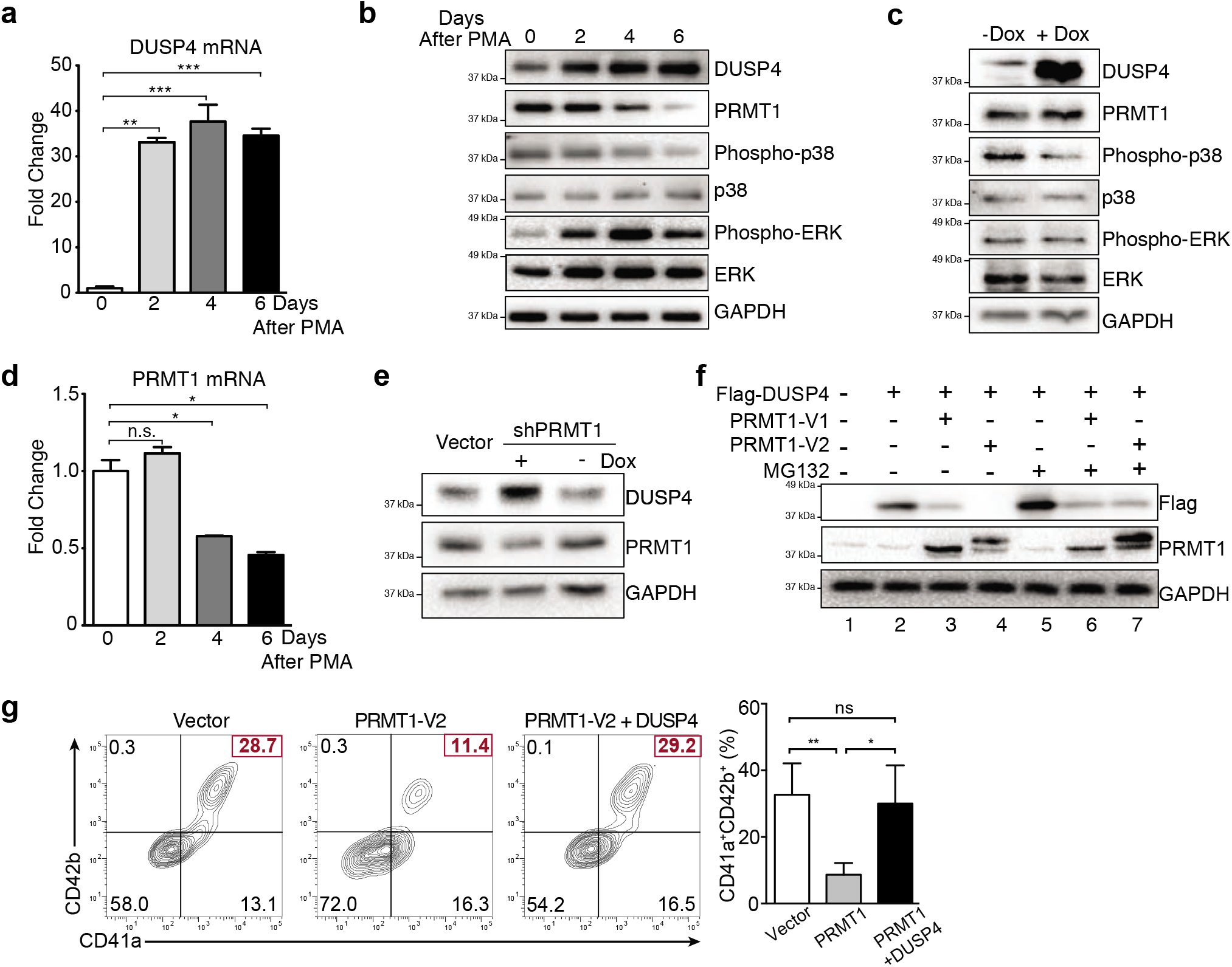
Crosstalk between DUSP4 and PRMT1 for MAPK signaling in Mk differentiation. **a.** DUSP4 mRNA level in PMA-treated MEG-01 cells. Cells were harvested at indicated time intervals and the extracted mRNAs were quantified by real-time PCR. **b.** MAPK-related proteins and PRMT1 in MEG-01 cells after PMA stimulation. Protein extracts were collected at indicated time intervals for western blotting. **c.** Regulation of MAPK signaling upon DUSP4 overexpression. MEG-01 cells were treated overnight with doxycycline to induce DUSP4 ectopically expressed from lentivirus. Cell extracts were collected for western blotting. **d.** PRMT1 mRNA level in MEG-01 cells during the course of PMA-stimulated Mk differentiation. **e** and **f.** PRMT1-dependent regulation of DUSP4 protein. NB4 cells that conditionally express shRNA against PRMT1 were treated with doxycycline to induce PRMT1 knockdown (**e.**). DUSP4- and PRMT1-encoding plasmids were transfected into HEK293T cells for their overexpression in the presence or absence of MG132 treatment (**f.**). **g.** Antagonistic roles of PRMT1 and DUSP4 on Mk differentiation of human CD34^+^ cells. Human CD34^+^ cells were infected with PRMT1 lentivirus (puromycin-R) and DUSP4 lentivirus (GFP), followed by puromycin selection and Mk differentiation. Percentages of CD41a^+^CD42b^+^ in GFP^+^ cells were determined by FACS. Representative plots (left panel) and statistics (right panel) were shown (n=3). Data were shown as mean ± SD, two-tailed paired t-test, *P ≤ 0.05; **P ≤ 0.01; ***P≤ 0.001.

DUSP4 has been shown to dephosphorylate MAPKs including ERK, JNK, and p38 kinases (37). To determine the specific MAPK(s) dephosphorylated by DUSP4, we performed Western blot analysis using lysates from PMA-treated MEG-01 cells to probe phosphorylation levels of ERK2, JNK, and p38 with the former two MAPKs implicated in Mk-Er differentiation. As cells underwent Mk differentiation, levels of phosphorylated p38 kinase decreased, while levels of phosphorylated ERK2 increased (**Fig. 3b**). Using a DUSP4-inducible cell line, we then confirmed that DUSP4 overexpression was sufficient for rapid suppression of p38 phosphorylation (**Fig. 3c**) but not the phosphorylation of ERK. This observation suggests that p38 signaling can be directly antagonized by high levels of DUSP4 to promote Mk differentiation. In contrast, JNK kinase phosphorylation levels were only modestly decreased during Mk differentiation (**Extended Data Fig. 2**). Therefore, the PRMT1-DUSP4 axis mainly suppresses the p38 kinase pathway for Mk differentiation. Furthermore, PRMT1-dependent activation of p38 kinase occurs in NB4 cells, which are immature granulocytes (**Extended Data Fig. 1e**). These consistent results in different types of cells imply that PRMT1-mediated p38 kinase activation can be achieved via direct suppression of DUSP4.

### PRMT1 controls DUSP4 stability by arginine methylation

We noted that the protein and mRNA expression levels of PRMT1 were inversely correlated with DUSP4 protein levels, but not with DUSP4 mRNA levels (**Fig. 3a, 3b** and **3d**). To determine whether PRMT1 activity may directly affect DUSP4 protein, we used doxycycline-inducible shRNA to knock down PRMT1, which resulted in increased DUSP4 protein levels (**Fig. 3e**). Conversely, overexpression of major PRMT1 isoform variants (V1 or V2) significantly decreased DUSP4 protein (**Fig. 3f**). An increased level of phosphorylated p38 kinase was detected in NB4 cells stably overexpressing PRMT1 (**Extended Data Fig. 1e**), which was associated with the decreased level of DUSP4. Treatment of PRMT1-overexpressing cells with the proteasome inhibitor MG132 blocked PRMT1-mediated DUSP4 degradation (**Fig. 3f**). These results collectively support that PRMT1 antagonizes DUSP4 protein levels by promoting proteasome-mediated target degradation. Of note, the PRMT1 isoform V2 exerts a stronger effect on protein degradation (compare Lane 3 and 4 in **Fig. 3f**).

To further elucidate the antagonistic relationship between DUSP4 and PRMT1 during Mk differentiation, we next examined the effects of DUSP4 on PRMT1-mediated block of Mk differentiation (**Fig. 3g**). Here, stimulation of human CD34^+^ hematopoietic progenitor cells with TPO and stem cell factor (SCF) toward CD41a^+^CD42b^+^ megakaryocytes was inhibited by PRMT1 overexpression, which decreased the frequency of Mk differentiation by three-fold (**Fig. 3f, Extended Data Fig. 3**). In contrast, expression of DUSP4 in PRMT1-overexpressing cells nearly restored the efficiency of megakaryocyte differentiation to levels comparable to CD34^+^ cells that do not overexpress PRMT1 (**Fig. 3g**). These results indicate an antagonistic interplay of the PRMT1-DUSP4 axis in regulating Mk differentiation into matured CD41a^+^CD42b^+^ cells.

### scRNA-seq analysis of Mk differentiation of primary human CD34^+^ cells

HSPCs (hematopoietic stem/progenitor cells) are heterogeneous with a continuum of transcriptional expression topology (8, 38). Novel regulatory pathways for Mk and Er differentiation have been discovered by scRNA-seq analysis (39, 40). In this context, we leveraged scRNA-seq technology in combination with FACS sorting to interrogate essential modulators of Mk differentiation in human CD34^+^ BM cells (**Fig. 4a**). A Drop-Seq protocol was implemented for scRNA-seq analysis of non-stimulated (day 0) and TPO/SCF-stimulated cells (day 8 post-stimulation). Total numbers of analyzed cells included 1,813 non-stimulated cells; 927 TPO/SCF-stimulated cells; 1,077 TPO/SCF-stimulated, CD41a^+^CD42b^+^ cells; and 867 TPO/SCF-stimulated, CD41a^-^ CD42b^-^ cells. The scRNA-seq data of all 4,684 cells were analyzed using the SPRING algorithm, a force-directed layout developed for visualizing continuous gene expression topologies and for assigning the cell fates of hematopoietic cells (**Fig. 4b**) (41). Unstimulated human CD34^+^ BM consisted of a mixture of cell subpopulations with lineage-specific gene expression signatures indicative of dendritic (D), granulocytic neutrophil (GN), lymphocytic (Ly), monocytic (M), and MPP cells (**Fig. 4c**). After 8 days of TPO/SCF stimulation, cell subsets characterized by lineage-specific gene expression signatures of basophilic cells (or mast cells), Er, Mk, and likely their progenitors (**Fig. 4c**), including a characteristic long “tail” on the SPRING plot that largely consisted of Mk cells. Analysis of the gene expression signature of CD41a^+^CD42b^+^ cells FACS-sorted from the TPO/SCF-stimulated cell population indicated that these cells overlapped with the tail population in the day-8 TPO/SCF-stimulated cell populations. Basophilic cells (or mast cells) were embedded within the TPO/SCF-stimulated, non-CD41a^+^CD42b^+^ cells, consistent with scRNA-seq analysis of HSPCs showing that basophils can be derived from MEPs (9, 40).

**Fig. 4.**
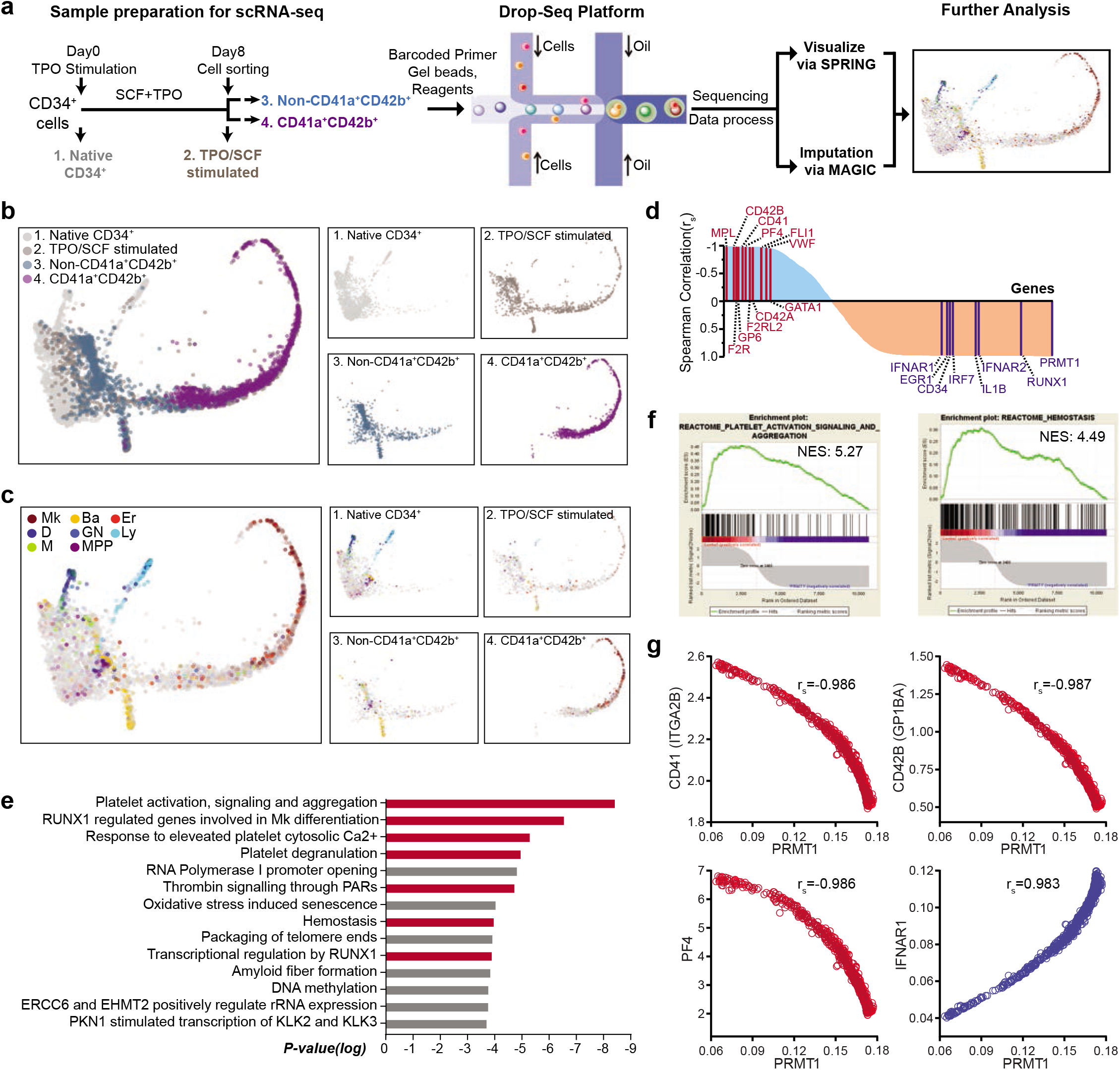
Negative correlation between PRMT1 and Mk differentiation revealed by single-cell RNA-seq (scRNA-seq) analysis. **a.** Experimental design for scRNA-seq analysis. Sample 1: Native CD34^+^ cells were cells isolated directly from bone marrow; Sample 2: CD34^+^ cells were cells cultured in TPO and SCF for 8 days before sorting with CD41a and CD42b; Sample 3: The non-CD41a^+^CD42b^+^ cells were sorted from sample 2; and sample 4 are CD41a^+^CD42b^+^ cells sorted from sample 2. **b.** and **c.** SPRING plots of single-cell transcriptomes. Individual cells were presented according to their origins (**b.**) or transcriptome-associated cell types (**c.**). Ba, basophilic or mast cell; D, dendritic; Er, erythroid; GN, granulocytic neutrophil; Ly, lymphocytic; M, monocytic; Mk, megakaryocytic; MPP, multipotential progenitors. **d.** Pathway analysis of the top 400 genes with the strongest negative Spearman correlation to PRMT1 expression level in the TPO/SCF-stimulated CD41a^+^CD42b^+^ population. Red bars highlight Mk-relevant biological pathways. **e.** Spearman correlation coefficients between PRMT1 and any gene revealed by scRNA-seq in TPO/SCF-stimulated CD41a^+^CD42b^+^ cells. Representative genes with significant correlation and functional relevance are annotated. **f.** Gene Set Enrichment Analysis (GSEA) of PRMT1-correlated genes in the TPO/SCF-stimulated CD41a^+^CD42b^+^ population. GSEA inputs are Spearman correlation coefficients of PRMT1 versus any gene with single-cell resolution. **g.** Normalized expression of representative transcripts co-plotted against that of PRMT1 transcript in TPO/SCF-stimulated CD41a^+^CD42b^+^ cells with single-cell resolution.

To further elucidate the TPO/SCF-stimulated CD41a^+^CD42b^+^ cell population, we implemented the MAGIC algorithm, a diffusion-based imputation method (42), to recover potential dropouts associated with inefficient mRNA capture (**Fig. 4a**). The resultant MAGIC-normalized scRNA-seq transcripts of the TPO/SCF-stimulated, CD41a^+^CD42b^+^ cells were analyzed using Spearman correlation between PRMT1 and other genes. The 400 transcripts with the largest and smallest Spearman correlation coefficients were then subject to pathway analysis with Reactome (**Fig. 4d** and **4e**, **Extended Data Table 1**). With the whole set of Spearman correlation coefficients as inputs, Gene Set Enrichment Analysis (GSEA) was also conducted to identify PRMT1-associated biological pathways (**Fig. 4f**) (43). Remarkably, the two approaches revealed Mk differentiation and platelet production, at single-cell resolution, as the top biological processes negatively correlated with the transcript level of PRMT1 (**Fig. 4e** and **4f**). Consistently, the transcript levels of CD41 (ITGA2B), CD42B (GP1BA), PF4, MPL, CD42A (GP9), GP6, VWF, FLI1, F2R, F2RL2, and GATA1, a collection of characteristic Mk differentiation markers (38, 44), negatively correlated with the transcript level of PRMT1 of individual cells (**Fig. 4e**, **4g** and **Extended Data Fig. 4**). In contrast, IFNAR1/2, EGR1, IRF7, and IL1B— a panel of proinflammatory response genes that can trigger p38 activation (45, 46) associated with blocking Mk differentiation (47, 48)—positively correlated with the transcript level of PRMT1 (**Fig. 4e, 4g** and **Extended Data Fig. 4**). We therefore conclude a negative correlation between PRMT1 and Mk-platelet differentiation at a single-cell resolution in the mature Mk cells as defined by CD41a^+^CD42b^+^ surface markers, and a positive correlation between PRMT1 upregulation and the genes associated with p38 kinase activation of proinflammatory pathways. These data support that as Mk cells mature, the PRMT1 expression level tapers off along with the decreased p38 kinase activity.

### Methylation of DUSP4 by PRMT1

Due to the antagonistic functions of PRMT1 and DUSP4 in Mk differentiation, we then examined whether DUSP4 is a substrate of PRMT1 in live cells. To profile PRMT1 methyltransferase activity, we developed the next-generation BPPM technology (**Fig. 5a** and **Extended Data Fig. 5a**) with the prior knowledge that the M48G variant of PRMT1 and the homologous M233G variant of PRMT3, but not wild-type PRMTs, utilized bulky SAM analogue cofactors to label their substrates with terminal-alkyne-containing chemical moieties (49). The three-step next-generation live-cell BPPM approach consisted of **(1)** the biosynthesis of *(E)*-hex-2-en-5-ynyl-SAM (Hey-SAM)—a bulky sulfonium-alky SAM analogue that is inert to wild-type methyltransferases—from its cell-permeable methionine analogue precursor Hey-Met by an engineered methionine adenosyl-transferase (MAT2A I117A) within live cells; (**2**) *in situ* modification of the PRMT1 substrates by the engineered PRMT1 M48G variant with Hey-SAM as a cofactor; and (**3**) enrichment and quantification of the Hey-SAM-modified substrates via the azide-alkyne click reaction with an azide-containing fluorescent dye (**Extended Data Fig. 5a** and **5b**) or biotin chemical reporter followed by enrichment with streptavidin-conjugated beads (**Fig. 5a**) (21, 50-52). The robust labeling of endogenous DUSP4 by PRMT1 (V1 or V2 isoforms), as well as the canonical substrate histone H4, was observed in HEK293T cells only in the presence of the complete set of the BPPM reagents---the engineered PRMT1, MAT2A, and *in situ* produced Hey-SAM---but not in the absence of either the engineered PRMT1 or the MAT2A variant (**Fig. 5b**). A similar labeling was also observed with exogenously expressed *N*Merminally-tagged DUSP4 variants in multiple cell lines, including Flag-tagged DUSP4 in HEK293T cells, and HA-tagged DUSP4 in K562 cells and NB4 cells (**Extended Data Fig. 5c** and **5d**). The next-generation live-cell BPPM technology thus demonstrated its ability to profile PRMT1 substrates within live cells. The consistent results of PRMT1-mediated DUSP4 methylation in these different cells demonstrate that DUSP4, like histone H4, can be robustly methylated by PRMT1 within multiple cellular settings.

**Fig. 5.**
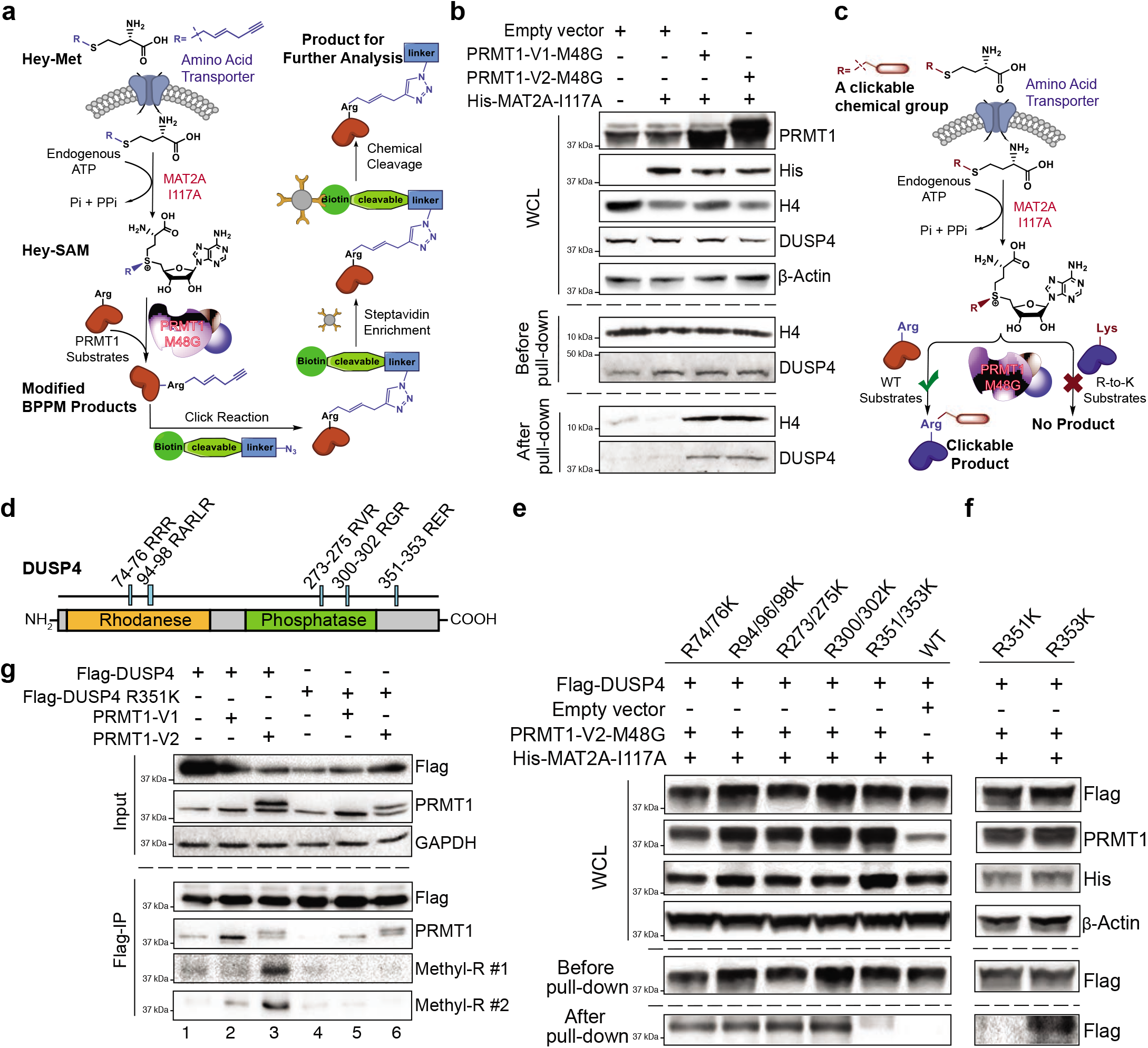
R351 methylation of DUSP4 by PRMT1. **a.** Schematic of the next-generation live-cell BPPM technology to uncover substrates of PRMT1. Cells that express PRMT1-M48G and MAT2A-I117A were treated with Hey-SAM, which is produced *in situ* from membrane-permeable Hey-Met inside live cells. PRMT1-M48G modifies PRMT1 substrates by Hey-SAM as a cofactor with its terminal-alkyne handle. The resulting modified proteins can be appended with a biotin tag via an azide-mediated click chemistry and then enriched by streptavidin beads for further analysis. **b.** Immunoblotting readouts of H4 and DUSP4 as PRMT1 targets enriched via the BPPM technology. HEK293T cells were treated with various combination of BPPM reagents. The resulting cell lysates were enriched by streptavidin beads followed by western blotting for the targets of interests. Samples before and after biotinstreptavidin pull-down were analyzed. **c.** Schematic description of the next-generation live-cell BPPM technology to reveal methylation site(s) of PRMT1. Candidates of methylated arginine residues on a protein substrate are mutated to lysine. The resulting Arg-to-Lys mutation is expected to diminish or abolish the BPPM-associated labeling. **d.** DUSP4 sequence with functional domains and RXR motifs highlighted. **e.** and **f.** Revealing PRMT1 methylation site(s) on DUSP4 with the next-generation live-cell BPPM technology. HEK293T cells were transfected with DUSP4 variants containing Arg-to-Lys mutation at putative sites. The BPPM-labeled products were enriched for quantitative comparison. The DUSP4 variants containing dual-Arg-to-Lys mutations at RXR motifs (**e.**) and the point mutations at R351 and R353 (**f.**), respectively. **g.** Validating PRMT1-invovled R351 methylation on DUSP4 with anti-methylarginine antibodies. HEK293T cells were transfected with PRMT1 and Flag-tagged DUSP4 (native or R351K mutant). Cellular extracts were immunoprecipitated with anti-Flag-antibody-conjugated resin. Inputs (upper panel) and pull-downs (bottom panel) were used for western blotting. Two anti-methyl-arginine antibodies were used to determine the presence of arginine methylation in DUSP4.

### PRMT1 methylates DUSP4 at R351

After identifying DUSP4 as a bona fide target of PRMT1 inside live cells, we modified the BPPM procedure to map the site(s) of PRMT1-mediated DUSP4 methylation (**Fig. 5c**). The potentially methylated Arg residues such as in the GAR motif (53) and arginine rich motif (20, 54-58) were mutated into Lys residues (**Fig. 5d**). Using the resulting R-to-K DUSP4 variants as substrate candidates of PRMT1, BPPM revealed that only the R351/352K, but not other mutations, significantly abolished the DUSP4 labeling activity of PRMT1 in HEK293T cells. This indicates that PRMT1 recognizes the RER motif (aa 351-353) of DUSP4 in live cells (**Fig. 5e**). With the individual R351K and R353K variants, we further defined that the R351K, but not R353K, mutation abolished the DUSP4 labeling activity of PRMT1 (**Fig. 5f**). A similar labeling pattern between the R351K variant and the R353K variant of DUSP4 was also observed in NB4 cells and K562 cells (**Extended Data Fig. 5c** and **5d**). Independently, PRMT1-dependent R351 methylation of DUSP4 was validated in cells expressing exogenous PRMT1 (V1 and V2 isoforms) and wild-type DUSP4, but not its R351K mutant, by two independently-developed pan-*anti*-methyl-Arg monoclonal antibodies from Cell Signaling Technology (**Fig. 5g**). Interestingly, the V2 isoform showed higher activity of DUSP4 methylation than the V1 isoform (**Fig. 5g**), consistent with the stronger effect of the former on DUSP4 degradation (**Fig. 3f**). Exogenously purified PRMT1, which robustly methylates histone H4 (59), also showed the ability to methylate the R351 residue of full-length DUSP4 as detected by mass spectrometry (**Extended Data Fig. 5e**). We also performed *in vitro* methylation assays using recombinant PRMT1 and (wild-type and R351A/F) DUSP4 proteins and further validated that DUSP4 is methylated by PRMT1 at the R351 site (**Extended Data Fig. 6**). These observations collectively demonstrate that PRMT1 predominantly methylates R351 of DUSP4 in biologically relevant contexts.

### PRMT1-mediated R351 methylation of DUSP4 promotes its degradation via poly-ubiquitylation

Since the methyltransferase activity of PRMT1 negatively regulates the stability of DUSP4 through proteasome-mediated degradation, a process expected to act via ubiquitylation (**Fig. 3f**), we further examined the antagonistic PRMT1-DUSP4 axis in the context of PRMT1-mediated R351 methylation of DUSP4. Consistent with PRMT1-promoted proteasome-mediated degradation of DUSP4, overexpression of PRMT1 (V1 and V2 isoforms) in HEK293T cells led to the robust poly-ubiquitylation of wild-type DUSP4 (**Fig. 6a**), which was completely suppressed on the DUSP4 R351K mutant that was inert for PRMT1 methylation (**Fig. 6a**). Furthermore, the stability of the DUSP4 R351K mutant was independent of the level of PRMT1, which contrasted with the decreased stability of wild-type DUSP4 upon PRMT1 expression (**Fig. 6b**). We assessed the stability of wild-type DUSP4 versus the methylation-dead R351K mutant in HEK293T cells (**Fig. 6c**). In these studies, wild-type DUSP4 showed a short half-life of less than 2 hours, while its R351K mutant was remarkably more stable, with an increased half-life of more than 5 hours (**Fig. 6c**). These data indicate that the R351 residue of DUSP4, and likely R351 methylation by PRMT1, are essential for promoting DUSP4 degradation (**Fig. 6d**). Consistent with these observations, the DUSP4 R351K variant was significantly more efficient than wild-type DUSP4 in promoting the Mk differentiation of human CD34^+^cells (**Fig. 6e**). Mass spectral analysis of immunoprecipitated DUSP4’s interactome revealed that HUWE1, one of the four E3 ligases associated with DUSP4, can account for the methylation-dependent degradation of DUSP4 as evidenced by HUWE1-DUSP4 interaction (**Fig. 6f**) and the promoted polyubiquitylation and then degradation of wild-type but not R351K DUSP4 (**Fig. 6g-i**). Overall, we conclude that the antagonistic role of PRMT1 on DUSP4 acts through a methylation-dependent, polyubiquitylation and proteome-mediated degradation of DUSP4. While this DUSP4-PRMT1 regulatory axis may occur in multiple cellular contexts, it plays an indispensable role in differentiation and maturation of Mk progenitor cells.

**Fig. 6.**
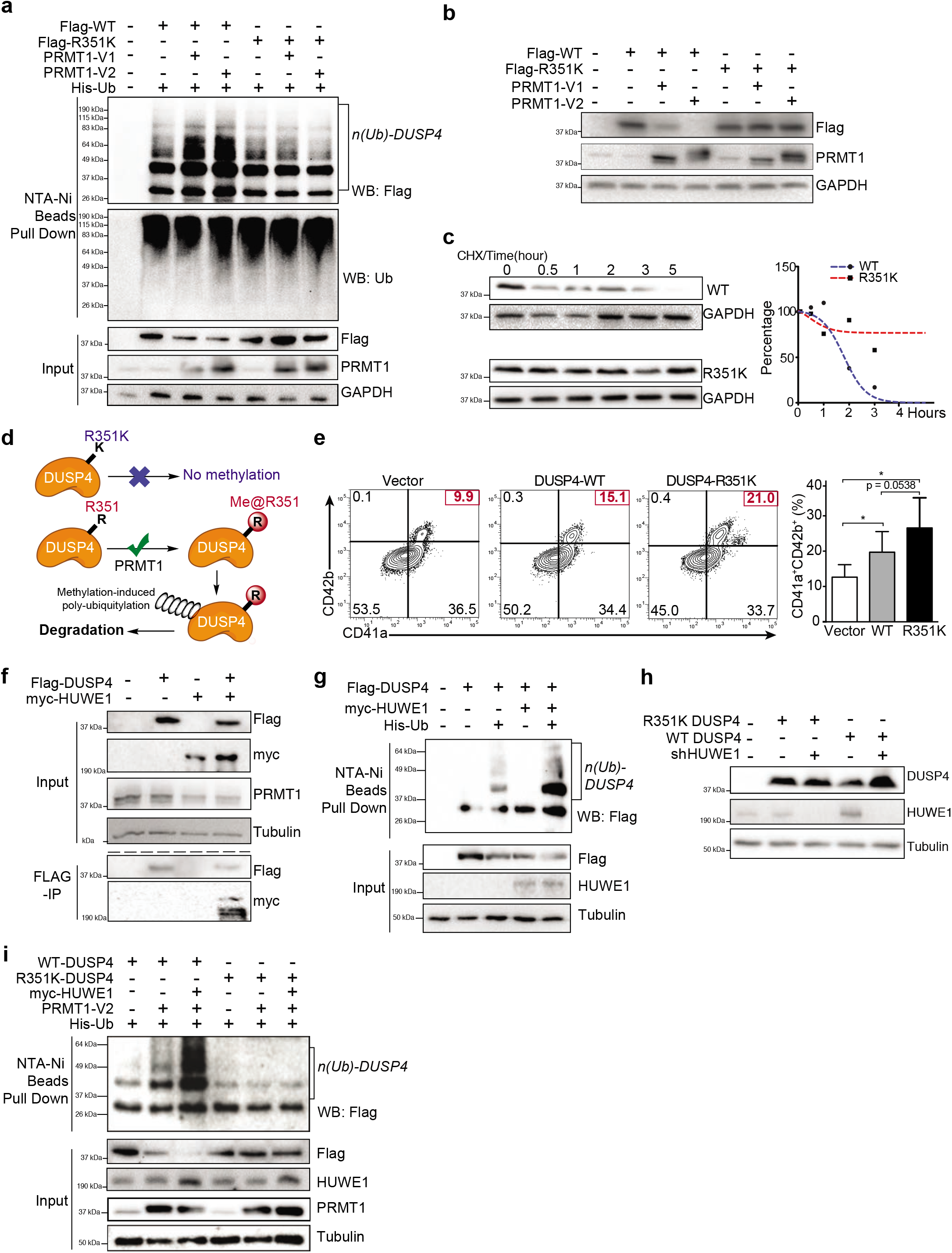
Poly-ubiquitylation and instability of DUSP4 promoted by PRMT1-involved R351 methylation. **a.** Poly-ubiquitylation of DUSP4, but not its R351K variant, stimulated by PRMT1. HEK293T cells were transfected with DUSP4 (wild-type and R351K mutant), 6×His-tagged ubiquitin and PRMT1 (V1 and V2 isoforms). Cells were treated with MG132 for 6 hours prior to harvest. NTA-Ni resin was used to isolate 6×His-ubiquitin-conjugated proteins. Fractions of pull-downs (upper panel) and inputs (bottom panel) were applied for western blotting. **b.** PRMT1-dependent stability of wild-type and R351K DUSP4. HEK293T cells were transfected with Flag-tagged DUSP4 (wild-type or R351K mutant) and PRMT1 (V1 and V2 isoforms). Cell extracts were harvested for western blotting. **c.** Half-life time of wild-type and R351K DUSP4. HEK293T cells were transfected with WT or R351K DUSP4, followed by the treatment of cycloheximide to block *de novo* protein synthesis. The resulting samples were harvested at indicated time intervals for western blotting. Normalized protein stability curves are plotted in the right panel. **d.** Mechanistic description of DUSP4 stability modulated by PRMT1-dependent R351 methylation. R351 methylation of DUSP4 by PRMT1 triggers its poly-ubiquitylation and thus its degradation; R351K mutation abolishes the methylation and thus suppresses poly-ubiquitylation and degradation. **e.** Mk differentiation in the presence of wild-type and R351K DUSP4. Human CD34^+^ cells were infected with lentiviruses expressing DUSP4s (wild-type or R351K mutant). The infected cells were cultured with SCF and TPO to induce Mk differentiation. Percentage of CD41a^+^CD42b^+^ cells was determined by FACS after 7 days. Representative plots and statistics were shown (n = 3). Data were shown as mean ± SD, two-tailed paired t-test, *P ≤ 0.05. **f.** Co-immunoprecipitation of HUWE1 and DUSP4. Flag-tagged DUSP4 was used to immunoprecipitate myc-tagged HUWE1 in co-transfected 293T cells. **g.** The protein ubiquitylation of DUSP4 is measured in 293T cells. Wild-type DUSP4 were expressed with or without myc-tagged HUWE1 and polyhistidine tagged ubiquitin. Western blots were performed with antibodies against Flag-tag,HUWE1 and tubulin as input controls and with antibody against Flag after the extract was affinity purified with NTA-Ni beads. **h.** Mutant and wild-type DUSP4 were expressed together with HUWE1 shRNA in 293T cells for western blotting with respective antibodies. **i.** Protein ubiquitylation assays with wild type and mutant DUSP4. DUSP4 wild type protein and mutant protein were expressed in 293T cells transfected with or without the plasmid combination of HUWE1 and PRMT1 as indicated on the top of the gel. The cell extracts were used for affinity purification with NTA-Ni beads to catch polyubiquitylated Flag tagged DUSP4 proteins.

### Pharmacological inhibition of p38 MAPK and PRMT1 stimulates Mk differentiation in myelodysplastic syndrome samples

To further explore the potential clinical application of the PRMT1-DUSP4 axis, we then characterized cells from patients with myelodysplastic syndromes (MDS), a myeloid malignancy characterized by low blood counts and defects in hematopoietic cell differentiation (60). Mk differentiation is adversely impacted in MDS, leading to thrombocytopenia and low platelet counts that are major clinical problems associated with this malignancy and that necessitate frequent platelet transfusion (61). Studies have shown that p38 MAPK is activated in small cohorts of MDS patients (62), which may suggest that the PRMT1-DUSP4 axis plays a role in the defective Mk differentiation. To address this question, we analyzed gene expression data of CD34^+^ HSPCs from MDS patients (N=183) and age-matched controls (N=17) (63). We found that PRMT1 levels were significantly elevated in MDS HSPCs (**Fig. 7a**). Furthermore, p38a MAPK was also significantly overexpressed in MDS HSPCs (**Fig. 7b**), which correlated with significantly lower platelet counts (**Fig. 7c**). Multivariate analysis using IPSS (international prognostic scoring system-the accepted system for prognosis in MDS) showed that p38a expression was predictive of adverse overall survival even after multivariate correction using standard risk factors as variables (**Fig. 7d**, Cox proportional hazards model, p = 0.049 for p38a expression using IPSS) (64). We further examined p38 phosphorylation status within megakaryocytes from 29 MDS patients using immunohistochemistry. Megakaryocytes from MDS patients had significantly higher levels of phosphorylated (activated) p38 kinase, in comparison with levels in normal control megakaryocytes (**Fig. 7e** and **7f**, **Extended Data Table 2** and **3**).

**Fig. 7.**
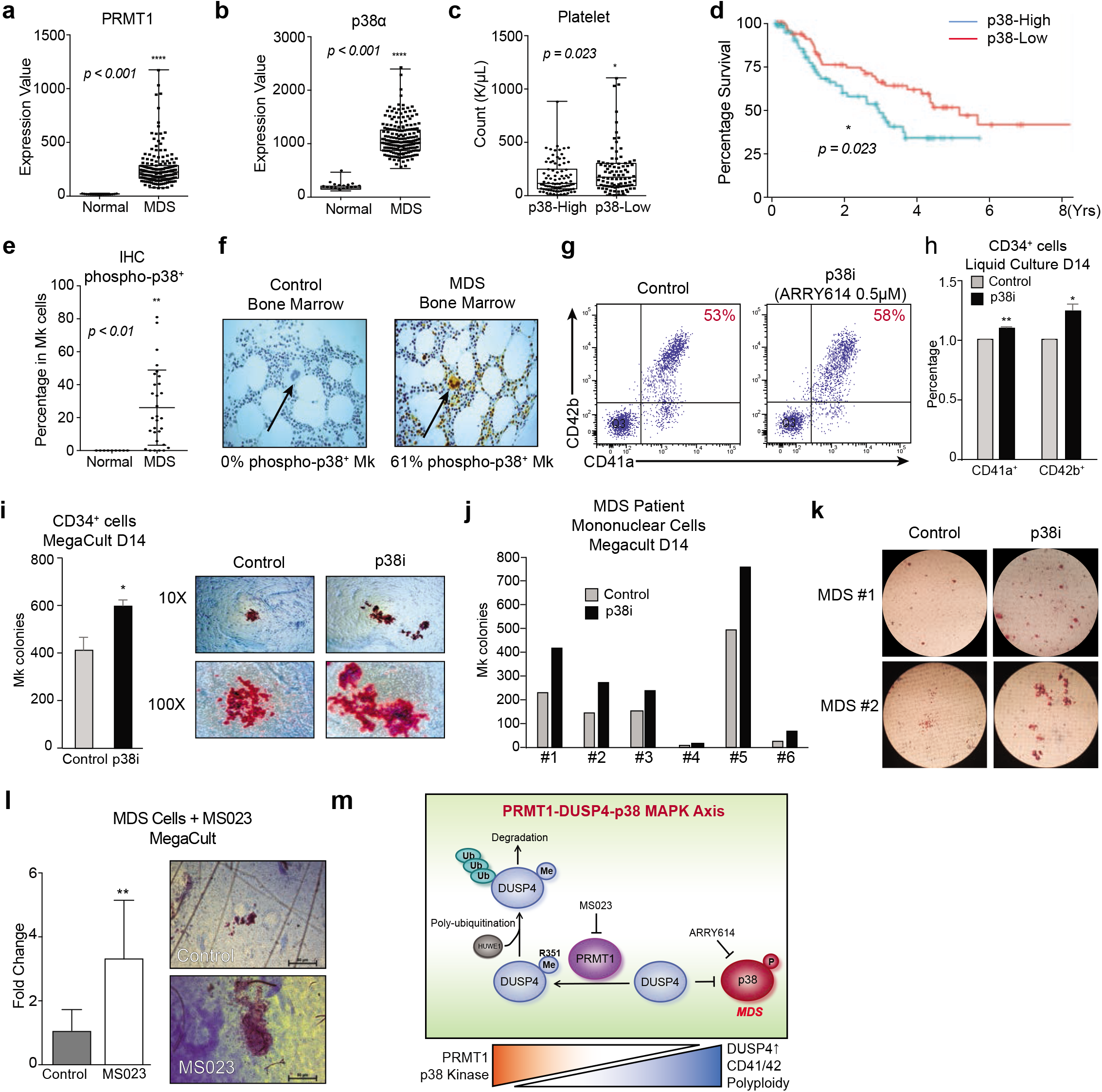
Clinical implication and pharmacological targeting of p38-DUSP4-PRMT1 axis in MDS. **a.** and **b.** Gene expression of PRMT1 and p38α in MDS patients and healthy donors with array-based analysis. The CD34^+^ HSPCs of MDS patients (N=183) and age-matched healthy controls (N=17) were analyzed for PRMT1 expression (**a.**) and p38α expression (**b.**). **c.** MDS cohorts as classified by low and high expression of p38α MAPK on the basis of median expression levels. The subjects with high p38α expression showed significantly lower platelet counts. **d.** Survival curves of MDS patients classified by low and high expression of p38α MAPK. The MDS patients with higher p38α expression in HSPCs showed significantly worse overall survival. **e.** and **f.** IHC analysis for phosphorylation-activated p38α MAPK of age-matched healthy controls and MDS bone marrow (BM) samples from a clinical trial with the p38α inhibitor Pexmetinib (ARRY614, labeled as “p38i”). MDS BMs showed significantly higher p-p38α staining in megakaryocytes (**e.**). Representative stains are shown (**f.**). **g. h.** and **i.** Effects of the p38α inhibitor Pexmetinib (labeled as “p38i”) on normal CD34^+^ cells. Normal CD34^+^ cells were grown in liquid culture conditions in the presence and absence of Pexmetinib and analyzed for CD41 and CD42 expression by representative charts shown (**g.**) and averaged FACS data (**h.**). Normal CD34^+^ cells were also grown for MegaCult assay for production of megakaryocyte colonies (**i.**). **j.** and **k.** Analysis of BM mononuclear cells (MNC) of MDS patients with MegaCult assay for production of megakaryocyte colonies in the presence or absence of the p38α inhibitor Pexmetinib (or “p38i”). Six MNC samples were examined (**j.**) with the representative images shown (**k.**). **l.** Analysis of MNC of MDS patients (n=4) with MegaCult assay for production of megakaryocyte colonies in the presence or absence of a PRMT1 inhibitor MS023. **m.** Mechanistic description of the signaling-epigenetic crosstalk during Mk differentiation via the PRMT1-DUSP4-p38 axis. Mk progenitors undergo abnormal differentiation in MDS by upregulation of PRMT1, which leads to p38 kinase activation. The relative levels of phospho-p38 are regulated by DUSP4. DUSP4 R351 is subject to PRMT1-medidated methylation, which leads to polyubiquitylation by HUWE1 and then degradation. Collectively, the PRMT1-DUSP4-p38 axis determines generation of Mk progenitor cells and the maturation of Mk cells.

A pilot Phase I clinical trial of the p38α MAPK inhibitor Pexmetinib (ARRY614) did not show significant increases in hemoglobin, but strikingly demonstrated increases in platelet counts in 12/16 (75%) of patients with transfusion-dependent severe thrombocytopenia (7/16 of patients achieved partial and 5/16 of patients achieved complete independence from platelet transfusion (ClinicalTrials.gov Identifier: NCT01496495) (65). For *ex vivo* validation of this striking observation, we examined terminal differentiation of human primary CD34^+^ cells in a liquid culture after p38 kinase inhibition (66). Pexmetinib treatment led to a modest but significant increase in Mk terminal differentiation (**Fig. 7g** and **7h**). Furthermore, in MegaCult assays that quantify MEP progenitor cells with the potential to differentiate to the Mk lineage, p38 kinase inhibition significantly promoted Mk colony formation (**Fig. 7i**). Importantly, when primary mononuclear cells from MDS patients were plated in the MegaCult assays after being treated *ex vivo* with Pexmetinib, we observed significant increases in Mk colonies for all six cases (**Fig. 7j** and **7k**). The enhancement of Mk differentiation with Pexmetinib treatment was also recapitulated using MS023 (67, 68), an inhibitor of type I PRMTs including PRMT1 (**Fig. 7l**). We tested MS023 inhibition of protein methylation in leukemia cell lines. MS023 at 200 nM inhibited more than half of the global protein arginine methylation as well as the methylation of two representative substrates RBM15 and DUSP4 (**Extended Data Fig. 7a** and **7b**). Furthermore, MS023 promoted Mk differentiation (more CD41a^+^CD42b^+^ population, **Extended Data Fig. 7c, 7d** and **7f**) and increased polyploidy (**Extended Data Fig. 7e**) in leukemia cells and primary human cells. Both the p38 kinase inhibitor and the PRMT1 inhibitor increased the colony sizes, which suggests that inhibition of the PRMT1-DUSP4-p38 pathway promotes the maturation of Mk cells derived from single Mk progenitor cells. These data indicate that pharmacological inhibition of p38 MAPK or PRMT1 may be a promising therapeutic strategy for treatment of thrombocytopenia in MDS patients. We also examined gene expression profiles generated from FACS sorted LT-HSCs (Lin^-^CD34^+^CD38^-^) and ST-HSC from 12 AML/MDS samples with normal karyotype, del(Chr7) and complex karyotype. We observed that HUWE1 was significantly overexpressed in leukemia stem cell populations (**Extended Data Fig. 8**).

## Discussion

### The next-generation live-cell BPPM technology reveals DUSP4 as a substrate of PRMT1

Our studies identified DUSP4 as a bona fide substrate of PRMT1 in multiple cellular settings using the next-generation live-cell BPPM technology (21, 50-52). In this context, the BPPM approach was leveraged using M48G PRMT1, a PRMT1 variant identified previously to effectively process bulky SAM analogs such as Hey-SAM while minimally acting on the native SAM cofactor (51). Hey-SAM, which is inert to native protein methyltransferases, contains a terminal-alkyne moiety that is transferred by the engineered PRMT1 onto PRMT1 substrates such as DUSP4. The resulting modified protein can then be labeled by an azide-containing biotin probe via the azide-alkyne click reaction and enriched by streptavidin-conjugated beads for further characterization. For live-cell BPPM, Hey-SAM is produced *in situ* by the I117A variant of MAT2A, which has been characterized previously to recognize broad *S*-alkyl methionine analogs as membrane-permeable substrates (50). With the livecell BPPM approach, quantitative comparisons can be made between native and Arg-mutated substrates, which showed that the R351 residue of DUSP4 is the dominant PRMT1 methylation site in multiple cellular contexts. While the BPPM technology was initially developed to interrogate methylome (a collection of methylation substrates) of designated protein methyltransferases, our work has adapted the technology to allow identification of individual substrate candidates and to reveal specific modification sites of PRMT1 in relevant cellular contexts. Given that the M48 residue of PRMT1 is conserved across the PRMT family and the homologous M233G variant of PRMT3 also utilizes bulky SAM analog cofactors (69), we envision that the next-generation live-cell BPPM approach will be generally applicable for profiling and validating PRMT targets.

To characterize proteome-wide Arg methylation, conventional approaches rely on mass spectroscopy (MS) or antibodies to trace characteristic methylarginine-containing peptides (49). However, because of the low abundance of many Arg methylation events, MS-based detection approaches often need to be coupled with antibody-based enrichment strategies. Further, methylation has minimal effects on the overall size and electrostatic property of the targeted Arg residues. The small difference in the biophysical properties between methylated and unmodified basic residues such as Lys and Arg thus makes it challenging to develop high-quality antibodies to probe methylation events in an unambiguous manner. This is particularly the case for Arg methylation of low abundance (24, 70). This situation is further complicated when Arg methylation is embedded in regions of the protein that are rich in other posttranslational modifications. Consequently, with conventional assays, many Arg methylation events and the associated PRMT activities are “invisible” in a cellular context. In contrast, BPPM technology allows effective enrichment of the targets of designated (engineered) PRMTs via distinct chemical labeling, whose detection is independent of neighboring sequences and posttranslational modifications. Additionally, the BPPM approach can unambiguously assign the peptides containing the distinctly labeled Arg residue(s) as the direct targets of a specific (engineered) PRMT activity. BPPM technology can also be coupled with conventional antibody-or MS-based assays to show that the BPPM-revealed Arg site(s) can be subject to native methylation (24, 70). Collectively, besides profiling the methylome of a designated PRMT, we further extended the utility of BPPM technology to validate specific PRMT substrates and identify the sites of the modification.

### Methylation of DUSP4 by PRMT1 promotes ubiquitylation

In this report, we identified that the R351 residue of DUSP4 is methylated by PRMT1 in multiple cellular contexts. The PRMT1 methylation site of DUSP4 (R351me) is in a SGPLRERGKTPAYP sequence, which also contains serine and threonine residues that have been shown to be phosphorylated (Phosphosite.org). Other RXRXXS/T-containing substrates of PRMT1 such as FOXO1 and BAD are methylated by PRMT1 on both Arg sites (56, 71), which blocks AKT-mediated phosphorylation on the nearby serine/threonine residue(s). The potential regulatory crosstalk between AKT phosphorylation and PRMT1 methylation of DUSP4 within this sequence remains to be studied. Notably, R351 is not conserved between DUSP1 and DUSP4. Although DUSP1 and DUSP4 share many biological pathways (72), upregulation of DUSP1 renders cancer cells resistant to chemotherapy and BCR-ABL kinase inhibitors (73, 74), while downregulation of DUSP4 enhances chemoresistance (75). Differential regulation via arginine methylation between DUSP1 and DUSP4 may distinguish their unique roles in hematopoiesis and in other developmental contexts.

PRMT1-mediated methylation of DUSP4 promotes its poly-ubiquitylation and proteasome-dependent degradation. Interestingly, methylarginine-promoted protein degradation has also been observed for other PRMT substrates such as SRC3 (76) and RBM15 (20). Consistently, we found that the PRMT1 isoform V2 more efficiently methylates DUSP4 than isoform V1 in cell lines, an effect that may be mediated by the extra amino acids on the N terminal region of V2. Since V2 may play a critical role in breast cancer cell proliferation (77, 78), understanding the structure and function of V2 warrants future study. One possible mechanism underlying methylarginine-promoted degradation is that an E3 ligase directly recognizes the methylarginine for binding. Alternatively, arginine methylation may directly trigger a change of protein conformation that enhances recognition by E3 ligases. Protein arginine methylation has been demonstrated to change protein conformation for liquid phase transition (79). Previously, we have shown that methylation of RBM15 by PRMT1 can trigger RBM15 ubiquitylation at a lysine site far from the methylation site (20). In addition, the E3 ligase responsible for RBM15 ubiquitylation is a RING domain E3 ligase CNOT4, while the E3 ligase responsible for DUSP4 is a HECT domain E3 ligase HUWE1. The two E3 ligases share no similar domains which can be predicted to recognize methylated arginines on their substrates. It will therefore be interesting to investigate whether arginine methylation is a commonly utilized signal for protein degradation via unconventional ways such as arginine methylated triggered liquid phase separation(79). Moreover, asymmetric di-methylated arginines (ADMA), which can only be generated from degradation of methylated proteins, cause high blood pressure (80). In this regard, upregulation of PRMT1 activity may have pathophysiological consequences, with ADMA as a secondary messenger for intercellular communication.

Here, we discovered that the E3 ligase HUWE1 is responsible for the methylation-dependent ubiquitylation and degradation of DUSP4. HUWE1 is a large protein containing multiple domains. A previous report demonstrated that HUWE1 recognizes phosphorylated protein for degradation (81), although the HUWE1 domains that recognize protein phosphorylation or methylation have not yet been identified. Furthermore, how HUWE1 expression levels impact PRMT1-regulated hematopoiesis and whether there is involvement of other E3 ligase(s) for the methylation-dependent degradation of DUSP4 remain to be investigated. Since PRMT1, DUSP4, and diverse E3 ligases are ubiquitously expressed in different tissues, the pathway may have distinct functions in different cellular contexts.

### Crosstalk between DUSP4 and MAPK signaling regulates hematopoietic differentiation

While DUSP4 can dephosphorylate ERK, p38, and JNK MAPKs *in vitro*, DUSP4 preferentially dephosphorylates ERK, JNK, or p38 in a highly cell-type-specific manner(82-84). Here, we identified a specific developmental context wherein DUSP4 fine-tunes MAPK signaling for determination of Mk cell fate at multiple differentiation steps (**Fig. 7m**). Given that the FACS sorted progenitor cells are still heterogeneous, we cannot examine whether DUSP4 determines the binary cell-fate choice at the level of a single MEP cell. Moreover, DUSP4 upregulation could happen at the stem-cell stage (85), and p38 activation mobilizes hematopoietic stem cells for differentiation(86, 87). Specifically, we showed that DUSP4 selectively dephosphorylates p38 (**Fig. 3**), while upregulation of ERK2 phosphorylation was not affected by DUSP4 expression during Mk maturation. Given that PRMT1 and DUSP4 are ubiquitously expressed, PRMT1-mediated methylation of DUSP4 may control MAPK signaling in other cellular processes. DUSP4 is a nuclear protein and DUSP4-mediated dephosphorylation of p38 kinase could happen on chromatin. The underlying DUSP4-MAPK signaling axis is expected to be coupled to other epigenetic events that make Mk differentiation irreversible. Signal transduction regulators such as p38 and ERK2 can be potential epigenetic regulators given their association with chromatin (88), and the intersection between the p38-DUSP4 and DUSP4-PRMT1 axis highlights the intrinsic connection between signaling events and epigenetic regulation (**Fig. 7m**).

DUSP4 knockdown in MEP cells specifically suppressed Mk colony-forming ability and promoted the generation of Er colonies (**Fig. 2**). As many stress signals can promote Er differentiation, the effects of DUSP4 on erythropoiesis could be indirect. In addition, MEP progenitor cells are very heterogeneous, so we cannot conclude that the binary switch really exists by turning on DUSP4 within specific time window of the MEP stage. Relatedly, at the erythrocyte maturation stage, excessive p38 kinase activity has been demonstrated to restrain excessive erythropoiesis by inducing apoptosis(89), suggesting that the role of p38 at different stages of Er differentiation could be different.

The scRNA-seq data of mature CD41a^+^CD42b^+^ cells demonstrate that p38 kinase activation is reduced during megakaryocyte maturation, while the p38-associated transcriptional signature such as interferon alpha receptor (IFNAR) is positively correlated with PRMT1 expression. Interestingly, inhibition of p38 kinase activity only modestly increased the number of CD41^+^CD42^+^ Mk cells in the liquid culture assay (**Fig. 7g**), which implies that the p38 kinase inhibitor may cause apoptosis along with differentiation during *in vitro* culture. On the other hand, p38 kinase inhibitor or PRMT1 inhibitor treatment yielded large size colonies (**Fig. 7g** and **7l**), suggesting that Mk progenitor cells in MDS patients regained the ability for maturation. This finding is further supported by the clinical observation that p38 kinase inhibitor reduces the need for platelet transfusion (65). We therefore expected that PRMT1 inhibitors enhance platelet production in MDS patients.

The activity of p38 kinase is downregulated during the maturation of CD41^+^CD42^+^ Mk cells, as demonstrated by the scRNA-seq data. Our data suggest that the later stage of Mk development and p38 MAPK activity are regulated by DUSP4, which also promotes Mk polyploidy (**Extended Data Fig. 1d**). Consistent with our data, it is reported that Mks with proinflammatory gene expression profiles are low polyploidy and reside in lung(90). Upregulation of PRMT1 may explain the inflammatory features observed in lung Mks. Data using the MDS patient samples further strengthen the model that inhibition of PRMT1 is sufficient for the formation of larger Mk colonies (**Fig. 7** and **Extended Data Fig. 7**). Our results and other studies have shown that PRMT1 expression and p38 MAPK activity are positively correlated with the interferon alpha response (**Fig. 4e**) (47), which is known to repress differentiation of megakaryocytes (91). These findings suggest that an additional role of DUSP4 in Mk differentiation could be to suppress expression of interferon response genes. Primary myelofibrosis (PMF) is a type of myeloid proliferative disease that is also characterized by abnormal megakaryopoiesis. P38 kinase is activated by FLT3 kinase in megakaryocytes of PMF (92). In PMF, inhibition of P38 kinase promotes Mk differentiation, and FLT3 kinase is directly methylated by PRMT1 for activation(93). Therefore, it would be interesting to further test PRMT1 inhibitors for PMF treatment.

### Anticipated benefit of inhibition of PRMT1 and p38 MAPK signaling for stimulating platelet production in MDS

Constitutive activation of p38 kinase in a subset of MDS cases (62) and its high expression can exhaust normal HSCs (86). Our data show that elevated p38 kinase activity and high levels of PRMT1 inhibit Mk differentiation, which may explain why MDS patients frequently develop thrombocytopenia. In light of the newly discovered PRMT1-DUSP4-p38 axis, we anticipated that suppression of overall p38 kinase activity in MDS cells using inhibitors such as Pexmetinib, or blocking PRMT1 activity with agents like MS023, could rescue Mk development and enhance platelet production. In a pilot phase I clinical trial, a large proportion of MDS patients with severe transfusion-dependent thrombocytopenia benefited from the p38α MAPK inhibitor, Pexmetinib, thus validating our preclinical findings. Our discovery of the inhibitory role of the PRMT1-DUSP4-p38 axis in megakaryopoiesis (**Fig. 7m**) provides a preclinical rationale for therapeutic targeting of components of this pathway in future clinical trials in patients with myeloid malignancies and thrombocytopenia.

