## Supplemental Information for "Methylation of Dual Specificity Phosphatase 4 Controls Cell Differentiation"

**Extended Information**

- **Material and Methods**
- **Reference for Methods**
- **Acknowledgements**
- **Author Contributions**
- **Statement of Conficting Interests**
- **Extended Data Figure Legends**

### Material and Methods:

**Human Primary CD34⁺ Cells**: The three sources of human primary CD34⁺ cells are peripheral blood, bone marrow and cord blood. The G-CSF-mobilized peripheral blood CD34+ cells were obtained in two steps: Ficoll-Paque Plus (GE Healthcare) to separate mononuclear cells and then CD34 MultiSort Kit (Miltenyi) for CD34 enrichment according to the description of the manufacturers. Human bone marrow and cord blood CD34+ cells were purchased from Lonza Inc. Human bone marrow samples were originated from a 35-year-old female and a 32-year-old male. The information of the donor(s) of cord blood is not available. These CD34+ cells were cultured in IMDM medium supplemented with 20% BIT (Stem Cell Technology) and a cytokine mixture consisting of 100 ng/mL SCF, 100 ng/mL FLT3 ligand, 50 ng/mL IL-6 and 20 ng/mL of TPO in 5% CO2 incubators at 37 °C before Mk stimulation. For viral transduction, these primary cells were spin-infected twice with corresponding lentiviruses (2 × 805 g for 2 hours with 8-hour interval).

**Cell Lines:** All of the used human cell lines were cultured in 5% CO2 incubators at 37 °C. HEK293T cells (human epithelia cell line derived from female embryonic kidney, RRID: CVCL_0063) were cultured in DMEM medium supplemented with 10% FBS. MEG-01 (human chronic myeloid leukemia cell line derived from a 55-year-old male, RRID: CVCL_0425), CMK (human megakaryocytic leukemia cell line derived from 10-year-old male with down syndrome, RRID: CVCL_0216), and NB4 (human acute promyelocytic cell line derived from a 23-year-old female, RRID: CVCL_0005) were cultured in RPMI1640 medium supplemented with 10% FBS. K562 (human chronic myeloid leukemia cell line derived from a 53-year-old female, RRID: CVCL_0004) cells were cultured in IMDM medium supplemented with 10% FBS.

**Bacteria:** BL21 (DE3) and Rosetta (DE3) cells were used for in vitro protein expression. These bacterial cells were cultured in LB medium at 37 °C.

**Antibodies:** anti-DUSP4/MKP2 antibody from Santa Cruz was used for the western blotting of BPPM experiments (**Fig. 5** and **Extended Data Fig. 3**), and anti-DUSP4 from Cell Signaling was used for all other western blotting (**Fig. 3** and **Extended Data Fig. 1**). For the staining of MDS cells, PE-CD41a and APC-CD42b antibodies (RRID: AB_396624, AB_398486) were used; for all other human CD34$^+$ cell staining, anther set of PE-CD41a and APC-CD42b antibodies (RRID: AB_395859, AB_398486) was used. See reagent list for detailed information for all antibodies.

**Cytokines:** for dual CFU-Mk/Er assays, cytokines used were purchased from Amgen and ConnStem. Cytokines used in other experiments were purchased from Peprotech.

**Plasmid constructs for shRNAs:** To screen the panel of DUSPs, shRNAs were designed using the SplashRNA algorithm [1] and expressed from SGEP (miR-E backbone) vectors as described previously [2]. The pLKO.1-shRNAs lentiviral vectors (DUSP4 #1 and #2) were purchased from Thermo Fisher. pLKO.1-GFP-shRNAs were created by replacing the puromycin-resistant element with the open reading frame of eGFP. See reagent list for detailed information.

**Plasmid constructs for gene expression:** (1) Transient transfection in HEK293T cells: cDNA of DUSP4 (WT or mutant) and PRMT1 (V1 or V2) were inserted into pCDNA.1 vector. (2) Differentiation assays of primary human CD34$^+$ cells: PRMT1 was inserted into pTripZ for Dox-inducible expression and DUSP4 was inserted into pBGJR-GFP for constitutive expression. (3) Stable cell line establishment: pBGJR-GFP-DUSP4 and pTripZ-DUSP4 were used for constitutive and inducible expression of DUSP4 in MEG-01 cells; pTripZ-PRMT1 was used for inducible expression of PRMT1 in NB4 cells. (4) BPPM experiments: pCDNA3.1 vector was used to express mutant versions of PRMT1, DUSP4 and MAT2A in HEK293T cells; pBGJR-GFP-PRMT1-M48G, pLV-EF1a-IRES-NEO-DUSP4 and pLEX-307-His MAT2A were used for protein expression in NB4 cells and K562 cells. See reagent list for detailed information.

**Synthesis of Hey-Met**: Hey-methionine analog (Hey-Met), *S*-(*E*)-hex-2-en-5-ynyl homocysteine, was synthesized by reacting *S*-adenosyl-L-homocysteine with (*E*)-6-bromohex-4-en-1-yne in the presence of AgClO4 in a 1:1 mixture of formic acid and acetic acid as described previously [3].

**Protein expression:** Native PRMT1 with N-terminal 6xHis tag was expressed in *E. Coli* Rosetta (DE3) Strain and purified by Ni-NTA agarose (Qiagen) and then HiTrap Q Sepharose Fast Flow Column (GE HealthCare) as described previously [4]. For DUSP4 expression, the plasmid harboring human DUSP4 with an N-terminal GST tag was transformed into BL21 (DE3) cells. The selected colony was cultured at 37 °C to $OD_{600}$ of 0.6~0.8. IPTG (0.1 mM) was added to induce protein expression for 4 hours. The cells were collected by centrifuging at 7,000 g at 4°C for 30 minutes. The resultant cell pellet was dissolved in a lysis buffer (50 mM Tris-Cl pH 8.0, 50 mM NaCl, 10% Glycerol and 0.1% Tween-20) supplemented with DNase I, lysozyme, and protease inhibitor cocktail (Roche), followed by French press for 4 times. This lysate was centrifuged at 30,000 g at 4°C for 1 hour. The supernatant was collected and applied to Glutathione Sepharose 4 Fast Flow GST-tagged protein purification resin (GE Healthcare) for DUSP4 purification. An elution buffer (50 mM Tris-Cl, pH 9.0, 50 mM NaCl, 10% Glycerol and 40 mM Glutathione) of 10 mL was added onto column, followed by ten collections of 1 mL fraction of eluates. To visualize and quantify DUSP4, Laemmli sample buffer was used to dissolve the samples. The resultant mixture was boiled and then resolved by SDS-PAGE. The gels were stained with Coommassie Brilliant Blue to determine the amount of DUSP4. Eluted fractions containing DUSP4 were combined and concentrated by centrifugation at 1,800 g at 4 °C with 30 kDa MWCO (Millipore). The concentrated DUSP4 were further purified by overnight dialysis in a dialysis buffer (50 mM Tris-HCl, pH 7.5, 100 mM NaCl and 15% Glycerol) at 4 °C with 20 kDa MWCO (Millipore).

**Mass Spectrometry:** Liquid chromatography-tandem mass spectrometry (LC-MS/MS) analysis was performed as described previously [5]. To prepare the Arg-methylated DUSP4 sample, 10 μM of GST-DUSP4 was incubated with 50 μM of $CH_3$-SAM and 2 μM PRMT1 in a reaction buffer (50 mM HEPES pH 8.0, 0.005% Tween-20, 0.0005% BSA and 1 mM TCEP) at ambient temperature (22 °C) overnight. The resulting reaction mixture was resolved by 4–12% Bis-Tris gel (Bio-Rad) and stained with Coomassie Brilliant Blue. The gel band corresponding to the methylated GST-DUSP4 was extracted and digested with trypsin (Promega) for LC-MS/MS analysis.

**Single-cell RNA Sequencing (scRNA-seq) Analysis:**

*Sample preparation.* Human bone marrow-derived $CD34^+$ cells were cultured in a pre-stimulation medium (IMDM + 20% BIT + Basic Cytokine Mix: 100 ng/mL SCF, 20 ng/mL TPO, 100 ng/mL FLT3 Ligand and 50 ng/mL IL-6) for four days and examined for the surface markers CD41a/CD42b before processing for scRNA-seq. For un-stimulated cells (Sample 1 in **Fig. 4**), approximately 100,000 cells were harvested. The rest of $CD34^+$ cells were cultured in a Mk-stimulating medium (IMDM + 20% BIT + MK Cytokine Mix: 25 ng/mL SCF, 50 ng/mL TPO), which was replaced with fresh Mk-stimulating medium on Day 4 and Day 7. On Day 8 post-stimulation, approximately 100,000 cells were collected as TPO/SCF-stimulated cells (Sample 2 in **Fig. 4**). FACS sorting of the TPO/SCF-stimulated cells allowed the collection of approximately 88,000 TPO/SCF-stimulated, $CD41a^+CD42b^+$ cells (Sample 4 in **Fig. 4**) and 100,000 TPO/SCF-stimulated, non-$CD41a^+CD42b^+$ cells (Sample 3 in **Fig. 4**). The four sets of cell samples were immediately subject to Drop-Seq-based single cell RNA sequencing as detailed below.

*Library preparation.* The procedure of scRNA-seq followed the C version 3.1 from Steve McCarroll's lab (http://mccarrolllab.org/wp-content/uploads/2015/05/Online-Dropseq-Protocol-v.-3.1-Dec-2015.pdf). Briefly, 1 mL of the samples containing approximately 100,000 cells with 90% viability were processed using microfluidic devices (PMDS Microfluidic Devices, FlowJEM). 8,000~10,000 cells were encapsulated in microfluidic droplets with 3000~4000 single cells recovered after washing and enzymatic steps. The resultant emulsion droplets were broken and subject to reverse transcription, followed by 14 cycles of PCR amplification---95 °C for 3 min, 4 x (98 °C for 20 s, 65 °C for 45 s, 72 °C for 3 min), 10 x (98 °C for 20 s, 67 °C for 20 s, 72 °C for 3 min), and 72 °C for 5 min. After purification with AMPure XP beads (Beckman Coulter), 600 pg barcoded cDNA was fragmented by Amplicon Tagment enzyme using Nextera XT Kit (Illumina), followed by indexing PCR--95 °C for 30 s, 12 x (95 °C for 10 s, 55 °C for 30 s, 72 °C for 30 s), and 72 °C for 5 min. The resulting DNA library was further purified with 0.6 x AMPure XP beads before sequencing.

*Sequencing and fastq generation.* Libraries were sequenced on Illumina NextSeq 500 platform (R1- 20 cycles, index1- 8 cycles, R2- 64 cycles). Samples were demultiplexed and raw fastq data were generated using bcl2fastq2 v2.19.

*Single-cell expression matrix generation.* Raw fastq data were processed and converted to single-cell expression counts matrix following the standard Drop-seq computational pipeline v1.12 developed by McCarrol lab (http://mccarrolllab.org/dropseq/). Briefly, barcodes and unique molecular identifiers (UMIs) were extracted and corrected for bead synthesis errors. Raw reads were trimmed for adapter and ploy-A sequences, followed by alignment to human GRCh37 reference genome using STAR v2.5.2b [6]. The number of captured cells was determined by selecting the “knee” point in the cumulative distribution of reads. The matrices of single-cell expression count were generated by counting UMIs per gene in captured cells.

*Single-cell data visualization with SPRING.* The cells detected with less than 100 genes or more than 4,000 genes or with > 20% mitochondrial RNA contents were excluded. After the sample filtering step, the scRNA-seq data of 1,813 cells, 1,077 cells, 864 cells, and 924 cells were obtained for Samples 1-4, respectively. The expression counts matrices on these filtered cells were uploaded to SPRING webserver (https://kleintools.hms.harvard.edu/tools/spring.html) for data visualization [7].

*Annotating cell types with signature genes.* Eight cell types were annotated with the current scRNA-seq data set using the algorithm code developed previously for mouse samples with minor modification (https://github.com/AllonKleinLab/klunctions/tree/master/sam/Paper_Code/Tusi_et_al_2018) [8]. Briefly, the lineage-specific mouse genes used in the original code were converted into human homologues: for E (Erythrocyte) with the mouse gene set of Hba-a2, Hba-a1, Alas2 and Bpgm replaced by the human gene set of HBA2, HBA1, ALAS2 and BPGM; for GN (granulocytes) with the mouse gene set of Lcn2, S100a8, Ltf, Lyz2 and S100a9 replaced by the human gene set of LCN2, S100A8, LTF, LYZ and S100A9; for Ly (Lymphocytic) with the mouse gene set of Cd79a, Vpreb3, Vpreb1 and Lef1 replaced by the human gene set of CD79A, VPREB3, VPREB1 and LEF1; for D (Dendritic) with the mouse gene set of Cd74, H2-Ab1 and Cst3 replaced by human gene set of CD74, HLA-DQB1 and CST3; for Meg (Megakaryocyte) with the mouse gene set of Pf4, Itga2b, Vwf, Pbx1 and Mef2c replaced by the human gene set of PF4, ITGA2B, VWF, PBX1 and MEF2C; for M (monocyte) with the mouse gene set of Csf1r and Ccr2 replaced by the human gene set of CSF1R and CCR2; for Ba (Basophilic) with the mouse gene set of Lmo4, Ifitm1, Ly6e and Srgn replaced by the human gene set of LMO4, IFITM1, LY6E and SRGN; for MPP (multipotent progenitor) with the mouse gene set of Hlf and Gcnt2 replaced by the human gene set of HLF and GCNT2.

*Recovering gene-gene expression relationship with MAGIC.* Certain gene-gene relationships in the current set of scRNA-seq data can be subject to random loss because of their low expressing and thus poor recovery. In order to overcome this technological limitation, MAGIC with default parameters [9] was applied to process the scRNA-seq data of the TPO/SCF-stimulated, $CD41a^{+}CD42b^{+}$ cells. Spearman correlations of PRMT1 transcript with other genes of individual TPO/SCF-stimulated, $CD41a^{+}CD42b^{+}$ cells were calculated with the following equation:

$$r_s = \frac{COV(r_{PRMT1}, r_x)}{\rho_{r_{PRMT1}} \rho_{r_x}}$$

Here, $\rho_{r_{PRMT1}}$ and $\rho_{r_x}$ are the standard deviations of the rank variables. $COV(r_{PRMT1}, r_x)$ is the covariance of the rank variables between PRMT1 and individual target genes.

*Pathway enrichment analysis.* Genes were ranked by spearman correlation coefficients. Reactome pathway enrichment analysis (https://reactome.org/) was performed using top 400 genes with positive and negative spearman correlation, respectively. Gene Set Enrichment Analysis (GSEA) analysis was performed using the full ranked list (10,656 genes) on curated gene sets (MSigDB, http://www.broadinstitute.org/msigdb).

**Virus Production and Stable Cell Line Establishment:** Viral constructs were co-transfected with envelope vectors and packaging vectors in HEK293T cells by the methods of calcium phosphate DNA transfection or lipofectamine 3000 (Invitrogen). psPAX2 and pMD2.G were used for lentiviral vectors; pCMV-VSVG and pCMV-dR8.2 were used for retroviral vectors. Harvested viral supernatants were used to spin-infect target cells. Stable cell lines were selected by either addition of 400 μg/mL G148, 2 μg/mL puromycin or by GFP-based flow cytometry sorting using BD FACSAria II system (BD).

**Bioorthogonal Profiling of Protein Methylation (BPPM):**

*Intracellular Labeling of PRMT1 Substrates.* The living-cell BPPM procedure described previously was followed with minor modification [3]. Briefly, constructs of PRMT1-M48G, His-tag MAT2A-I117A and Flag-tag DUSP4 (WT or R351K) were transfected into HEK293T cells. At 24 hours post transfection, cells were washed with PBS and replenished with fresh DMEM medium. At 48 hours post transfection, the old medium was removed and cells were incubated with a methionine-deficient medium (DMEM without methionine, glutamine and cysteine, supplemented with 10% dialyzed FBS, L-glutamine at 0.584 g/L and L-cystine.2HCl 0.0626 g/L) for 30 minutes. 0.5 mM Hey-Met was then added into the medium for 8-hour incubation before harvest. Alternatively, the cells that stably expressed PRMT1-M48G, MAT2A or DUSP4 were directly incubated in the methionine-deficient medium for 30 minutes before Hey-Met addition.

*Cell Harvest and Lysate Preparation.* After intracellular labeling with Hey-SAM processed from Hey-Met, cells were harvested, washed with PBS and then lysed in HEPES buffer (50 mM HEPES pH 8.0, 0.05% Tween-20, 1 mM TCEP, and Roche protease inhibitor cocktail) for 20 minutes on ice, followed by 20 minutes of sonication. Sample was centrifuged at 13,400 g at 4 °C for 30 minutes and the resulting supernatant was saved as lysates. Protein concentrations of the lysates were determined by Bradford assay and the expression of PRMT1, MAT2 or DUSP4 was verified by western blotting. Equal amounts of proteins (10~15 mg) were applied to methanol precipitation at -80 °C overnight as described previously [3].

*Click Reaction of Hey-SAM-labeled proteins with Biotin-containing Azide Probes.* The click reaction procedure described previously was followed with minor modification [10]. Briefly, after the step of methanol precipitation, the protein pellets (usually 5~10 mg) containing Hey-SAM-labeled PRMT1 substrates were resuspended in 1 mL of the resuspension buffer consisting of 50 mM Triethanolamine pH 7.4, 150 mM NaCl, 4% SDS and the Roche protease inhibitor cocktail. The sample was diluted up to 3.45 mL with 50 mM Triethanolamine (pH 7.4). Click reaction cocktail was prepared by mixing 200 µL of fresh 20 mM $CuSO_4$ and 200 µL of 40 mM BTTP (Click Chemistry Tools) for 30-60 minutes. 50 µL of freshly prepared 200 mM sodium ascorbate was added to the pre-mixed $CuSO_4$/BTTP, as the blue color disappears, followed by adding 100 µL of 10 mM diazo biotin-azide (Click chemistry Tools) DMSO stock solution. 550 µL of the click reaction cocktail was then added to the resuspension of protein lysates to reach the final condition: 1 mM $CuSO_4$, 2 mM BTTP ligand, 2.5 mM sodium ascorbate and

250 µM diazo biotin-azide. The sample was incubated at ambient temperature (22 °C) for 2 hours with shaking. Pre-chilled methanol was added to the click reaction mixture for overnight protein precipitation at -80 °C, followed by centrifugation at 3,200 g at 4 °C for 30 minutes. Protein pellets were washed twice with pre-chilled methanol then air-dried for further analysis.

*Biotin-Streptavidin pull-down.* Biotin-labeled proteins were then subject for Streptavindin pull-down as described previously with minor modification [10]. Protein pellets obtained from the click reaction of the diazo biotin-azide probe were resuspended in 1 mL of resuspension buffer containing 50 mM Triethanolamine pH 7.4, 150 mM NaCl, and 4% SDS with Roche EDTA-free protease inhibitors. The samples were resuspended to the equal final concentrations of the total proteins (1~5 mg per mL) determined by Bradford assay. Equal amounts of proteins (1~5 mg per mL) were then transferred into new tubes containing 2 mL of the dilution buffer consisting 50 mM Triethanolamine pH 7.4, 150 mM NaCl and 1% Brij97. Pierce High Capacity Streptavidin Agarose beads (Thermo) were pre-equilibrated with PBS for three times and then resuspended with the dilution buffer. The diluted protein samples were mixed with 200 µL of streptavidin beads and incubated for 1 hour at ambient temperature (22 °C) with end-over rotation. The beads were thoroughly washed with PBS containing 0.2% SDS (volume%) once, with PBS twice and 250 mM ammonium bicarbonate twice. During each wash, the samples were centrifuged at 3, 200 g for 2 minutes at 4 °C. The washed beads were transferred to spin columns (Amicon Ultra-0.5 ml Centrifugal Filter). Labeled proteins were cleaved from the beads by the treatment of 250 µL of the elution buffer consisting of 100 mM freshly prepared $Na_2S_2O_4$, in a buffer containing 250 mM Ammonium bicarbonate and 1% SDS for 30 minutes at ambient temperature (22 °C) with shaking. Eluted proteins were collected by centrifuging at 1,000 g at 4 °C for 2 minutes. The elution step was repeated once. The combined eluates were then transferred to 3 NMWL Centricon filter device (Millipore) to remove recessive $Na_2S_2O_4$. The protein samples were concentrated through lyophilization and then dissolved in laemmli loading buffer for further analysis.

*Click Reaction of Hey-SAM-labeled proteins with Florescent Azide Probes for In-gel Analysis.* Hey-SAM-labeled proteins were subject to click reaction with an azide dye for in-gel fluorescence visualization as described previously with minor modification [10]. After the step of methanol precipitation of Hey-SAM-labeled PRMT1 substrates as describe above, the protein pellets (5~10 mg) were resuspended in 1 mL of the resuspension buffer consisting of 50 mM Triethanolamine pH 7.4, 150 mM NaCl, 4% SDS and the Roche protease inhibitor cocktail. For in-gel fluorescence analysis, around 400 μg aliquant of individual protein samples was diluted up to 4-5 mg/mL with the resuspension buffer mentioned above. The 25 μL aliquant of this protein mixture was diluted up to 85 μL with 50 mM Triethanolamine (pH 7.4). Click reaction cocktail was prepared by mixing 5 μL of 20 mM freshly prepared $CuSO_4$ and 5 μL of 40 mM BTTP for 30-60 minutes. 2.5 μL of freshly prepared 200 mM sodium ascorbate was added to the pre-mixed $CuSO_4$/BTTP, as the blue color disappears, followed by adding 2.5 μL of 10 mM TAMRA-Azide (Thermo Fisher Scientific) DMSO stock solution. 15 μL of the click reaction cocktail was then added to the resuspension of protein lysates. The sample was incubated at ambient temperature (22 °C) for 2 hours with shaking. Pre-chilled methanol was added to the click reaction mixture for overnight protein precipitation at -80 °C, followed by centrifugation at 3,200 g at 4 °C for 30 minutes. Air-dried protein pellets were resuspended in a loading buffer consisting of 40 mM Tris pH=6.8, 70 mM SDS, 10 mM EDTA, 10% glycerol and 10% β-ME (without dye to avoid interference with fluorescence signal) and resolved by SDS-PAGE. The gels were then fixed in mixture of 40% methanol and 10% acetic acid (volume%) overnight to remove unreacted TAMRA dye. The gels were then rinsed in $H_2O$ for rehydration and scanned for fluorescence signal using the TAMRA channel on a Typhoon TRIO variable mode imager (Amersham Bioscience). The scanned gel was then stained with Coomassie Blue to examine the protein loading.

**Western Blotting:** Laemmli sample buffer was added to cell lysates and enriched proteins. Samples were boiled, sonicated (WCL), resolved by SDS-PAGE, and then transferred to PVDF membranes (Millipore) or nitrocellulose membranes (Bio-Rad). Membranes were blocked in 5% non-fat milk, followed by blotting of indicated primary and secondary antibodies. Luminata Western Chemiluminescent reagent (Millipore) and Bio-Rad ChemiDoc MP system (Bio-Rad) were used for final visualization and quantification.

**FLAG-Immunoprecipitation:** Calcium phosphate transfected HEK293T cells ($10^7$cells) were lysed in 1 mL H-Lysis buffer (20 mM HEPES pH 7.9, 150 mM NaCl, 1 mM $MgCl_2$, 0.5% NP40, 10 mM NaF, 0.2 mM $NaVO_4$, 10 mM β-glycerol phosphate and 5% glycerol) containing freshly prepared dithiothreitol/DTT (1 mM), PMSF

(200 μM) and the Roche protease inhibitor cocktail. Cells were incubated on ice for 15~30 minutes and then sonicated by Bioruptor Ultra-sonication system (Diagenode). The sonicated cell extracts were cleared by centrifugation at 12,000 g at 4 °C for 10 minutes and then applied for immunoprecipitations. 40 μL of pre-equilibrated 50% anti-FLAG M2 agarose (SIGMA) was added to samples, followed by overnight rotation at 4 °C. Precipitates were collected by centrifugation at 135 g at 4 °C for 2 minutes and sequentially washed twice by each of three buffers: (1) 25 mM HEPES pH 7.9 and 150 mM NaCl; (2) PBS + 0.2% Triton X-100; and (3) PBS. Hot 1x Laemmli sample buffer was added to washed beads, followed by 5-minute boiling to release bound proteins.

Ubiquitylation Assay. HEK293T cells were transfected with the DUSP4 (WT and mutant) construct and the construct of His-ubiquitin as previously described with minor modification [11]. Briefly, MG132 was added to the culture at 40 hours post transfection and kept in the culture for 6 hours before harvest. Approximately $10^7$ cells were lysed in 1 mL of buffer A (6 M guanidine-HCl, 0.1 M $Na_2HPO_4$/ $NaH_2PO_4$ pH 7.5, 10 mM imidazole) on ice for 10 minutes. Lysate was briefly sonicated (< 5 cycles) using Bioruptor (Diagenode). Lysates were centrifuged at 12,000 g for 10 minutes at 4 °C. Supernatants were collected and incubated with 100 μL of pre-balanced $Ni^{2+}$-NTA beads (QIAGEN) for 3 hours at ambient temperature with occasional mixing. The NTA beads were collected by centrifugation and washed with buffer A, buffer B (1.5 M guanidine-HCl, 25 mM $Na_2HPO_4$/ $NaH_2PO_4$, 20m M Tris-Cl pH 6.8, 17.5 mM imidazole) and buffer TI (25 mM Tris-Cl pH 6.8, 10 mM imidazole). After the final washing, the beads were resuspended in hot 1x Laemmli sample buffer supplemented with 200mM imidazole. Samples were boiled and then used for western blotting.

**Half-life Assay:** HEK293T cells were used for transfection of DUSP4 constructs (WT or R351K). At 36 hours post transfection, cells were supplemented with fresh medium containing 25 μg/mL cycloheximide (CHX). Extracts were collected at 0, 0.5, 1, 2, 3 and 5 hours post CHX treatment. Laemmli sample buffer was added to extracts. Samples were boiled and used for western blotting.

**MDS cells in vitro Mk Colony Formation Assays:** For characterization of Mk colonies (CFU-Mk), MegaCult-C CFU-Mk was used. Patient bone marrow- or peripheral blood- derived mononuclear cells (10,000 cells/mL) were used for each assay and incubated for 14 days before colony quantification. MS023 (500 nM) and ARRY-614 (200 nM) were added into culture in respective assays.

**Immunohistochemistry:** Paraffin-mounted bone marrow core biopsy sections were collected from MDS patients or control subjects with consent. Bone marrow section staining was performed with anti-p-p38 antibody (Cell signaling, Cat #9211) as previously described[12].

**In Vitro Arginine Methylation Assay:** The *in vitro* methylation of DUSP4 was measured by a previously reported filter plate assay with some modifications[10, 13]. The reaction mixture (a total volume of 20 µL) contains 100 nM SETD8 protein (*N*-terminal 6 × His tagged), 1 µg DUSP4 protein (*N*-terminal GST-tagged), 0.75 µM [$^3$H-Me]-SAM (PerkinElmer Life Sciences), and reaction buffer (50 mM HEPES, pH = 8.0, 0.005% BSA, 1 mM TCEP). The resulting mixture was allowed to react at ambient temperature (22°C) overnight. Each reaction mixture was split into three aliquots and quenched by spotting on phosphor cellulose P-81) filter paper, followed by 2 hour air-dry. The dried filter paper was then washed 5 times with 50 mM $Na_2CO_3$/$NaHCO_3$ solution (pH = 9.2). The washed filter paper was then transferred into a scintillation vial, well mixed with 0.5 ml dd$H_2O$ and 5 ml Ultima Gold, and analyzed by a Liquid Scintillation Analyzer (Perkin Elmer Tri-Carb 2910 TR).

**MDS Patient Sample Gene Expression Dataset and Survival Analysis:** Gene expression data from 183 MDS CD34$^+$ samples and 17 controls were obtained from Gene Expression Omnibus (GEO) (GSE19429)[14]. Survival analysis and blood counts were correlated with expression of p38 MAPK and PRMT5 as done previously [15]

**Mk Differentiation of Primary CD34+ Cells:** For screening DUSPs relevant to Mk differentiation, lentiviruses encoding shRNAs against individual DUSP transcripts were used to infect CD34$^+$ cells. The transfected cells were stimulated with 50 ng/mL TPO and 25 ng/mL SCF for eight days before FACS analysis of GFP/CD41a$^+$/CD42b$^+$ population on BD Accuri™ C6 (BD Biosciences). To further validate Mk differentiation perturbed by DUSP4 shRNAs, lentivirus-infected CD34$^+$ cells were cultured in IMDM supplemented with 20% BIT, 50 ng/mL TPO and 2 ng/mL SCF for seven days. Expression of CD41a and CD42b were determined by flow cytometry on BD LSRFortessa (BD Biosciences). To examine binary Mk/Er differentiation, CD34$^+$ cells were culture in IMDM with supplement of 20% BIT, 50 ng/mL TPO, 2 ng/mL of SCF and 5 U/mL EPO for ten days. Expression of CD41a and CD71 were determined by flow cytometry on BD LSRFortessa (BD Biosciences).

**Mk Differentiation of MEG-01 Cells:** MEG-01 cells or lentivirus-infected MEG-01 cells were cultured in RPMI1640 with 10% FBS. 20 nM PMA (phorbol myristate acetate) were added to the medium to promote Mk differentiation. Expression of CD41a was determined by flow cytometry. Propidium iodide (PI) staining was used to assess the total DNA content. The cell population with >4N DNA was characterized as polyploid.

**Dual CFU-Mk/Er Assay:** The populations of MEP were purified from parental or lentivirus-infected $CD34^+$ cells via FACS-sorting as previously described [16]. MEP were sorted with $Lin^-CD34^+FLT3^-CD45RA^-MPL^+CD38^{mid}CD41a$. For each assay, 500 sorted cells were plated on MegaCult™-C Medium with lipids and the supplement of TPO (50ng/mL), EPO (3U/mL), IL-3 (10ng/mL), IL-6 (10ng/mL) and SCF (25ng/mL). At 14 days of culturing, colonies were fixed and stained with anti-GlyA and anti-CD41a. Colonies were counted and categorized based on the stain of GlyA and CD41a: CFU-Mk ($GlyA^-CD41a^+$), BFU-E ($GlyA^+CD41a^-$) and CFU-Mk/E ($GlyA^+CD41a^+$).

**Real-time PCR:** Total RNA was prepared using Direct-Zol RNAprep Kit (Zymo research). cDNA was generated by the Verso cDNA Synthesis Kit (Thermo Scientific) with random hexamer primers. Real-time PCR assays were performed with Absolute Blue qPCR SYBR green Mix (Thermo Scientific) on a ViiA 7 system (Applied Biosystems).

**Staining intracellular phospho-p38 for FACS analysis:** Approximately $1x10^6$ cells of NB4 cells were collected, washed twice with cold PBS, and fixed in 0.4% PFA for 7 minutes on ice. The cells were then washed with cold PBS and fully fixed in 4% PFA for 15 minutes at ambient temperature (22 °C). The fixed cells were washed twice with FACS Buffer (PBS + 2% BSA + 0.1% $NaN_3$) and then permeabilized with methanol for 30 minutes at -20 °C. The resultant cells were washed with the FACS buffer two more times before staining with antibodies for 45 minutes at ambient temperature (22 °C). The antibody-stained cells were then washed with the FACS buffer twice before FACS analysis.

**Quantification and statistical analysis:** The 2-tailed Student's t test was used for significance testing in the bar graphs. *P value*s less than 0.05 were considered significant. Quantification data are presented as mean ± SEM.

**Data Availability:** Single Cell RNA seuquencing dataset will be available at online archive. All other data is available in the main text or extended information.

**Meterial List:**

| REAGENTS OR RESOURCES | SOURCE | IDENTIFIER |
|---|---|---|
| **Antibodies** | | |
| anti-6x His tag | Abcam | Cat# 18184; RRID: AB_444306 |
| anti-actin | Abcam | Cat# 3280; RRID: AB_303668 |
| anti-DiMethyl-R #1 | Guo et. al. Mol Cell Proteomics, 2014[17]; AMDA-D4H5 | Not commercially available |
| anti-DiMethyl-R #2 | Guo et. al. Mol Cell Proteomics, 2014[17]; AMDA-D6H8 | Not commercially available |
| anti-DUSP4 | Cell Signaling | Cat# 5149; RRID:AB_2750867 |
| anti-Erk | Cell Signaling | Cat# 9102, RRID:AB_330744 |
| anti-FLAG | Sigma-Aldrich | Cat# F3165; RRID:AB_259529 |
| anti-GAPDH | Thermo | Cat# MA5-15738; RRID:AB_10977387 |
| anti-CD41a | BD Bioscience | Cat# 555465; RRID:AB_395857 |
| anti-CD235a | Bio-Rad | Cat# MCA506; RRID:AB_323506 |
| anti-H4 | Cell Signaling | Cat# 13919; RRID: AB_2798345 |
| anti-HA Tag | Millipore | Cat# 05-904; RRID: AB_11213751 |
| anti-Mouse HRP | Santa Cruz | Cat# SC-2314; RRID: AB_641170 |
| anti-MKP2 | Santa Cruz | Cat# SC-1200; RRID: AB_2095314 |
| anti-p38 | Santa Cruz | Cat# SC-81621, RRID:AB_1127392 |
| anti-Phospho-Erk | Santa Cruz | Cat# SC-7976, RRID:AB_2297323 |
| anti-Phospho-p38 | Santa Cruz | Cat# SC-17852-R, RRID:AB_2139810 |
| anti-PRMT1 | Millipore | Cat # 07-404; RRID: AB_310588 |
| anti-Rabbit HRP | Santa Cruz | Cat# SC-2004; RRID: AB_631746 |
| anti-Tubulin | Proteintech | Cat# 66031-1; RRID: AB_11042766 |

| | | |
|---|---|---|
| anti-Ubiquitin | Sigma-Aldrich | Cat# 07-2130; RRID:AB_11205591 |
| APC Mouse anti-human CD42b | eBioscience | Cat# 17-0429-42, RRID: AB_2573146 |
| APC Mouse anti-human CD42b | BD Bioscience | Cat# 551061; RRID:AB_398486 |
| APC Mouse anti-human CD71 | BD Bioscience | Cat# 55137; RRID:AB_3985004 |
| ImmPRESS Anti-Rabbit Ig Reagent antibody | Vecotr Laboratories | Cat# MP-7401; RRID:AB_2336529 |
| PE Mouse anti-human CD41a | BD Bioscience | Cat# 557297; RRID:AB_396624 |
| PE Mouse anti-human CD41a | BD Bioscience | Cat# 555467; RRID:AB_395859 |
| PE Phospho-p38 MAPK | Invitrogen | Cat# 12-9078-42; RRID:AB_2572691 |
| PE Phospho-pS6 Ribosome Protein | Cell Signaling | Cat# 5316S; RRID:AB_10694989 |
| PerCP/Cy5.5 anti-human CD41 | Biolegend | Cat# 303720; RRID: AB_2561732 |
| **Bateria and Virus Strains** | | |
| One Shot™ BL21(DE3) | Invitrogen | Cat# C6000033 |
| **Biological Samples** | | |
| Human bone marrow CD34+ progenitor cells | Lonza | 2M-101B |
| Human cord blood CD34+ progenitor cells | Lonza | 2C-101A |
| **Chemicals, Peptides, and Recombinant Proteins** | | |
| [3H-Me]-SAM | PerkinElmer Life Sciences | Cat# NET155V |
| 4–12% Bis-Tris gel | Bio-Rad | Cat#3450124 |
| Ammonium bicarbonate | Sigma-Aldrich | Cat# 9830 |
| anti-FLAG M2 agarose | Sigma-Aldrich | Cat# A2220 |
| BIT | Stem Cell Technology | Cat3 09500 |
| BTTP ligand | Chemical Synthesis Core, Albert Einstein College of Medicine | Not Applicable |
| cOmplete Protease Inhibitor Cocktail | Roche | Cat# 11697498001 |
| Copper(II) sulfate pentahydrate | Sigma-Aldrich | Cat# 203165 |
| Cycloheximide | Cayman | Cat# 14126 |
| Dialyzed Fetal Bovine Serum | Gibco | Cat# 26400-044 |
| Diazo Biotin-Azide | Click Chemistry Tools | Cat# 1041-25 |
| DMEM without glutamine, methionine, and cystine | Gibco | Cat# 21013024 |
| DMSO | Sigma-Aldrich | Cat# D8418 |
| Dnase I | Roche | Cat# 10104159001 |
| DPBS | Corning | Cat# 21-030-CV |

| | | |
|---|---|---|
| Dulbecco's Modified Eagles Medium (DMEM) | GE Healthcare | Cat# SH30243.01 |
| Fetal Bovine Serum | GE Healthcare | Cat# SH30910.03 |
| Glutathione Sepharose 4 Fast Flow GST-tagged protein purification resin | GE Healthcare | Cat# 17513202 |
| Human EPO | Amgen | |
| Human EPO | Peprotech | Cat# 100-64 |
| Human FLT3L | Peprotech | Cat# 300-19 |
| Human IL3 | ConnStem | Cat# I 1003-A |
| Human IL3 | Peprotech | Cat# 200-03 |
| Human IL6 | ConnStem | Cat# I 1006 |
| Human IL6 | Peprotech | Cat# 200-06 |
| Human Stem Cell Factor | ConnStem | Cat# S 1000 |
| Human Stem Cell Factor | Peprotech | Cat# 300-07 |
| Human TPO | ConnStem | Cat# T 1002 |
| Human TPO | Peprotech | Cat# 300-18 |
| Iscove's Modified Dulbecco's Medium (IMDM) | GE Healthcare | Cat# SH30228.01 |
| Lipofectamine 3000 | Invitrogen | Cat# L3000008 |
| Luminata Western Chemiluminescent reagent | Millipore | Cat# WBLUC0500 |
| Lysozyme | MP Biochemicals | Cat# 100834 |
| MG132 | Cayman | Cat# 10012628 |
| $Ni^{2+}$-NTA beads | QIAGEN | Cat# 1018244 |
| Nitrocellulose Membrane | Bio-Rad | Cat# 1620167 |
| Pierce™ High Capacity Streptavidin Agarose beads | Thermo | Cat# 20359 |
| PMA | Cayman | Cat# 10008014 |
| PVDF Membrane | Millipore | Cat# PFL00010 |
| RPMI 1640 | Corning | Cat# 10-040-CV |
| RPMI 1640 without methionine | Gibco | Cat# A1451701 |
| SAM (S-Adenosyl-Methionine) | Sigma-Aldrich | Cat# A7007 |
| Sodium hydrosulfite | Sigma-Aldrich | Cat# 157953 |
| Triethanolamine | Sigma-Aldrich | Cat# 90279 |
| Trypsin | Promega | Cat# V5280 |
| **Commercial Assay Kits** | | |
| CellTiter-Glo Viability Assay Kit | Promega | Cat# G7572 |
| Direct-Zol RNAprep Kit | ZYMO | Cat# R2062 |

| | | |
|---|---|---|
| ImmPACT Vector Red Substrate Kit | Vecotr Laboratories | Cat# SK-5105; RRID:AB_2336524 |
| ImmPRESS™ HRP Anti-Mouse IgG Polymer Detection Kit | Vecotr Laboratories | Cat# MP-7452; RRID:AB_2744550 |
| MegaCult™-C Medium with Lipids | STEMCELL | Cat# 04850 |
| MegaCult™-C Complete Kit with Cytokines; Collagen | STEMCELL | Cat# 04901, 04902 |
| Methylcellulose Base Media kit | R&D | Cat# HSC002 |
| Verso cDNA synthesis Kit | Thermo | Cat# AB1453 |
| **Experimental Models: Cell Lines** | | |
| CMK | DSMZ | Cat# ACC-392; RRID: CVCL_0216 |
| HEK293T | ATCC | Cat# CRL-3216; RRID: CVCL_0063 |
| K562 | ATCC | Cat# CCL-243; RRID: CVCL_0004 |
| MEG-01 | ATCC | Cat# CRL-2021; RRID: CVCL_0005 |
| NB4 | Dr. Stephen Nimer | RRID: CVCL_0005 |
| **Oligonucleotide** | | |
| **list of primers** | | |
| PRMT1-Fwd | 5' CCA GTG GAG AAG GTG GAC AT | |
| PRMT1-Rev | 5' CTC CCA CCA GTG GAT CTT GT | |
| DUSP4-Fwd | 5' AGG CGG CTA TGA GAG GTT TT | |
| DUSP4-Rev | 5' CAC TGC CGA GGT AGA GGA AG | |
| HPRT1-Fwd | 5' CAC CCT TTC CAA ATC CTC AG | |
| HPRT1-Rev | 5' CTC CGT TAT GGC GAC CCG CA | |
| **sGEP shRNA Sequences** | | |
| Control | shRen.713 | |
| DUSP1-a | TTATGTAACAAAATGTCTTCTT | |
| DUSP1-b | TTGTATAAAAAAGTCATCCTTA | |
| DUSP2-a | TCTGTATAAATATAAAGTGCTA | |
| DUSP2-b | TTCTGTATAAATATAAAGTGCT | |
| DUSP4-a | TACAACAACGACAACAAAGGGA | |
| DUSP4-b | TATTTCTAGAGGAAGCAGGGAG | |
| DUSP5-a | TTAAAAACAAAAATCACAGTTG | |
| DUSP5-b | TTTTCTTCACATTCACACGGGT | |
| DUSP6-a | TTAGTATTAACCAATTCCGCAC | |
| DUSP6-b | TAAAGAGAAAAAATCATGAGGA | |
| DUSP7-a | TTGAGTGACAGGTTCATCTTCT | |
| DUSP7-b | TAAAAAGTAAACTCAGGTCCGT | |
| DUSP8-a | ATAATATACATTTATAACGGGC | |

| | | |
|---|---|---|
| DUSP8-b | TATAATATACATTTATAACGGG | |
| DUSP9-a | AAGAAAGCAACTATGATATGGA | |
| DUSP9-b | TAAATAAACTGTTTATTCAGAA | |
| DUSP10-a | TAAACAAAGGTTGAGATCCTGA | |
| DUSP10-b | TTAATTTGTCAGTTTGTGGGAG | |
| DUSP16-a | TATATTTCAGATTTACAGGGAA | |
| DUSP16-b | ATATATTTCAGATTTACAGGGA | |
| **pLKO.1 shRNAs sequences** | | |
| Control | Scramble | |
| DUSP4-#1 | GCCTACCTGATGATGAAGAAA | |
| DUSP4-#2 | CCCAGTGGAAGATAACCACAA | |
| HUWE1 | AAACCCAGGGCTGCCTTGGAAAAG | |
| **Recombinant DNA** | | |
| Plasmid: pcDNA3.1 | Thermo Fisher | Cat# V79020; Addgene: #2093 |
| Plasmid: pGEX-6p-1 | GE HealthCare | Cat# 28-9546-48; Addgene: #2887 |
| Plasmid: pLKO.1 | David Root | Addgene: #10878 |
| Plasmid: pLEX_307 | David Root | Addgene: #41392 |
| Plasmid: pBGJR-GFP | Dr. Vladimir Jankovic | Not commercially available |
| Plasmid: pLV-EF1a-IRES-Neo | Hayer et al. Nat Cell Biol. 2016[18] | Addgene: #85139 |
| Plasmid: pTripZ | Thermo Fisher | Addgene: #5561 |
| **Software and Algorithms** | | |
| STAR | Dobin et al., 2013[6] | https://github.com/alexdobin/STAR |
| SPRING | Weinreb et al., 2018[7] | https://kleintools.hms.harvard.edu/tools/spring.html |
| MAGIC | van Dijk et al., 2018[9] | https://www.krishnaswamylab.org/projects/magic |
| **Other** | | |
| BD Accuri C6 | BD Biosciences | N/A |
| BD FACSAria II | BD Biosciences | N/A |
| BD LSRFortessa Special Order System | BD Biosciences | N/A |
| Bio-Rad ChemiDoc MP system | Bio-Rad | Cat# 170-8280 |
| Bioruptor Ultra-sonication system | Diagenode | Cat# UCD-300 |
| PDMS Microfluidic Devices | FlowJEM | N/A |
| Synergy H1 plate reader | BioTek | Cat# 8041000 |
| TRI-CARB 4910TR 110 V Liquid Scintillation Counter | PerkinElmer Life Sciences | Cat# A491000 |

**Acknowledgments**

This study was supported by grants from the National Institutes of Health of USA (1R21CA202390 and 1R01DK110574 to XZ; R35GM131858 to ML; R01GM122749 to JJ; R01GM126154 to YGZ), National Cancer Institute of USA (5P30 CA008748 to ML via MSKCC), the Tri-Institutional PhD Program in Chemical Biology (HG), and US Department of Veterans Affairs (BX0003617 and BX004426 to YC). We would like to thank Dr. Iannis Aifantis at NYU for HUWE1 reagents.

**Author Contribution**

H.S., M.J., C.S., S.A., H.G., H.D., A.V., M.L., and X.Z. designed the study. H.S, M.J., C.S., S.A., J.X.F., N.T.T., S.M.L., H.G., S.J., Y.Z., C.W.S., X.H., and Y.C. performed the experiments. Q.Z., L.C., Y.S., J.L., J.J., and Y.G.Z. provided key reagents. M.J., T.Z., J.X., and M.L. performed computational analysis. C.K.Q., C.A.K., R.B., Y.C., S.D.N., H.D., D.S.K., J.X., A.V., M.L., and X.Z. supervised the project. H.S., M.J., C.S., A.V., M.L., and X.Z. prepared the manuscript with help from all coauthors.

### Declaration of Competing Interests

M.L. has served as a member of the Scientific Advisory Board for Epi One Inc.. A.V. has received research funding from GlaxoSmithKline, Incyte, MedPacto, Novartis, Curis, and Eli Lilly and Company; has received compensation as a scientific advisor to Novartis, Stelexis Therapeutics, Acceleron Pharma, and Celgene; and has equity ownership in Stelexis Therapeutics. Other authors have no conflicts of interest relevant to this work. Array BioPharma provided the p38 inhibitor Pexmetinib (ARRY614) and participated in its Phase I study (ClinicalTrials.gov Identifier: NCT01496495).

### Extended Data Figure Legends

**Extended Data Fig. 1. Requirement of DUSP4 for optimal Mk differentiation of MEG-01 cells.**

**a.** Heatmap of the percentages of CD41a$^+$ cells upon knockdown of DUSPs in MEG-01 cells. MEG-01 cells were infected with lentiviruses containing a control RNA or shRNAs against DUSPs. Two shRNAs were used for each DUSP with a replicate. Percentages of CD41a$^+$ cells in each DUSP knockdown were normalized to those of the paired control. Normalized ratios are indicated in the heatmap in color scale.

**b.** and **c.** Representative FACS and rigorous statistical analysis are respectively shown for the data above with 3 independent replicates (n = 3).

**d.** Analysis of Mk differentiation of MEG-01 cells upon DUSP4 overexpression. MEG-01 cells and MEG-01-DUSP4 cells were collected for FACS analysis of CD41a and DNA content (PI staining). Cells with >4N of DNA content were characterized as polyploid.

**e.** FACS analysis of the intracellular phospho-p38 level of parental and PRMT1-overexpressing NB4 cells. Cells were fixed by 0.4% PFA and then 4% PFA, and permeabilized by methanol for antibody staining. Negative and positive controls for NB4 cells without antibody stain and stained with *anti*-PE-phospho-S6K control antibody, respectively. Cells were harvested at indicated time points for western blotting. Data were shown as mean ± SD, two-tailed paired t-test, *$P \leq 0.05$; **$P \leq 0.01$.

**Extended Data Fig. 2. JNK kinase phosphorylation was modestly changed during Mk differentiation of MEG01 cells.**

**a.** MEG-01 cells were stimulated with PMA. The cell lysates were harvested on the indicated days for western blotting with antibodies shown the left side of blots.

**b.** The ratio of band densities of activated kinases to total kinase levels.

**Extended Data Fig. 3. Verification of overexpression of DUSP4 and PRMT1 in virus-transduced CD34$^+$ cells.**

**a.** Verification of gene expression after viral transduction. CD34$^+$ cells were cultured in basic cytokine mix medium (IMDM, 20%BIT, SCF, FLT3L, TPO and IL-6) for two days before viral infection. Viral vectors pBGJR-DUSP4 and pTripZ-PRMT1V2 were used to express genes, and empty virus were used as controls. Three rounds of infection were performed with 12-hour gaps. Forty-eight hours post the last infection, puromycin was added to culture to select pTripZ infected cells. After 72 hours of selection, GFP+ cells (pBGJR-infected) were collected by flow cytometry. Expression of transduced genes were verified by qPCR.

**b.** Whole-cell extract of sorted cells were used for western blotting.

**Extended Data Fig. 4. Single-cell RNA sequencing analysis of the Spearman correlation between PRMT1 and the transcript signature associated with Mk differentiation.**

**a.** and **b.** Normalized transcripts of Mk-relevant genes negatively and positively correlated with PRMT1 in Mk-stimulated CD41a$^+$CD42b$^+$ cells with single-cell resolution.

**Extended Data Fig. 5. Further validation of DUSP4 arginine methylation by BPPM and mass spectrometry.**

**a.** Schematic description of the BPPM technology coupled with in-gel fluorescence visualization. BPPM-labeled substrates were conjugated with TAMRA-dye via a click reaction, followed by electrophoresis and in-gel visualization.

**b.** BPPM-revealed substrates of PRMT1 with in-gel fluorescence as readout. HEK293T cells were transfected with various combination of BPPM reagents. Their expression was confirmed by immunoblotting. The cell lysate was subject to TAMRA-dye via the click reaction, followed by in-gel fluorescence visualization. The total protein loading was shown by Coomassie Brilliant Blue staining.

**c.** and **d.** Revealing substrate(s) and methylation site(s) of PRMT1 with the next-generation live-cell BPPM technology. NB4 cells (**c.**) and K562 cells (**d.**) exposed with various combinations of BPPM reagents and DUSP4 (wild-type and R351K mutant) were treated with membrane-permeable Hey-Met, which produces the SAM analogue cofactor Hey-SAM *in situ*, as described in Fig. 5. Modified substrates were enriched by streptavidin beads and subject to western blotting. Samples before and after biotin-streptavidin pull-down were analyzed.

**e.** Mass spectrometric analysis of methylated DUSP4 peptides. Recombinant DUSP4 was incubated with recombinant PRMT1 and SAM followed by SDS-PAGE separation. The DUSP4 band was digested with trypsin and subject to mass spectrum analysis. The fragment pattern was assigned as a DUSP4 peptide containing R351 methylation.

**Extended Data Fig. 6. *In vitro* methylation of DUSP4 by PRMT1**. Recombinant PRMT1 was used to methylate recombinant wild-type and mutant DUSP4 protein. The PRMT1 protein and DUSP4 proteins were purified using Ni-bead via histidine tags. Scintillation counts of different reactions were performed by incubating recombinant PRMT1 and DUSP4 proteins with $H^3$-SAM.

**Extended Data Fig. 7. PRMT1 inhibitor MS023 blocks arginine methylation and enhances Mk differentiation.**

**a.** Arginine methylation level of MS023-treated cells. MEG01 and NB4 cells were cultured with the presence of control or MS023 for 24 hours. Cell extract was harvested and used for SDS−PAGE. Arginine methylation status was detected by using anti-methyl-R antibody.

**b.** Inhibition of DUSP4 methylation by MS023. Flag-tagged DUSP4 was immunoprecipitated from 293T cells treated with MS023. Two generic anti-methyl-arginine antibodies were used for western blotting.

**c.** CMK cells were treated with MS023 for 72 hours then stained with anti-CD41a/anti-CD42b antibodies for FACS analysis.

**d.** Histograms of MS023-treated CMK cells labeled with anti-CD41a and anti-CD42b antibodies, respectively.

**e.** Polyploidy of MS023-treated CMK cells were assessed by propidium iodide staining. Cells with DNA content of >4N were counted as polyploidy.

**f.** Immobilized peripheral CD34+ cells were cultured in Mk-promoting medium (TPO + SCF) with or without the presence of MS023. Cultured cells were harvested on day 7 for FACS analysis.

**Extended Data Fig. 8. HUWE1 gene expression level is high in AML/MDS stem cells.**

Analysis of the HUWE1 gene expression levels in LT-HSC and ST-HSC cells

Su and Jiang et.al. Extended Data Fig. 1

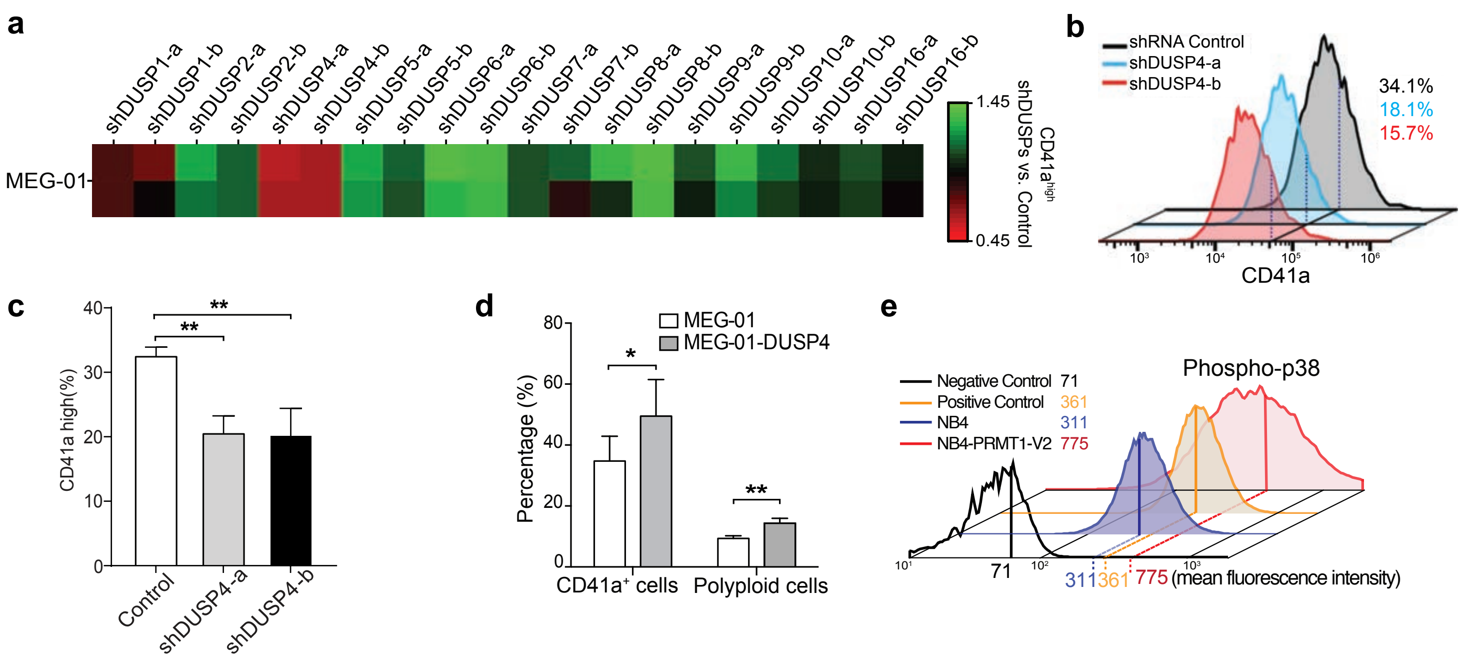

Su and Jiang et.al. Extended Data Fig. 2

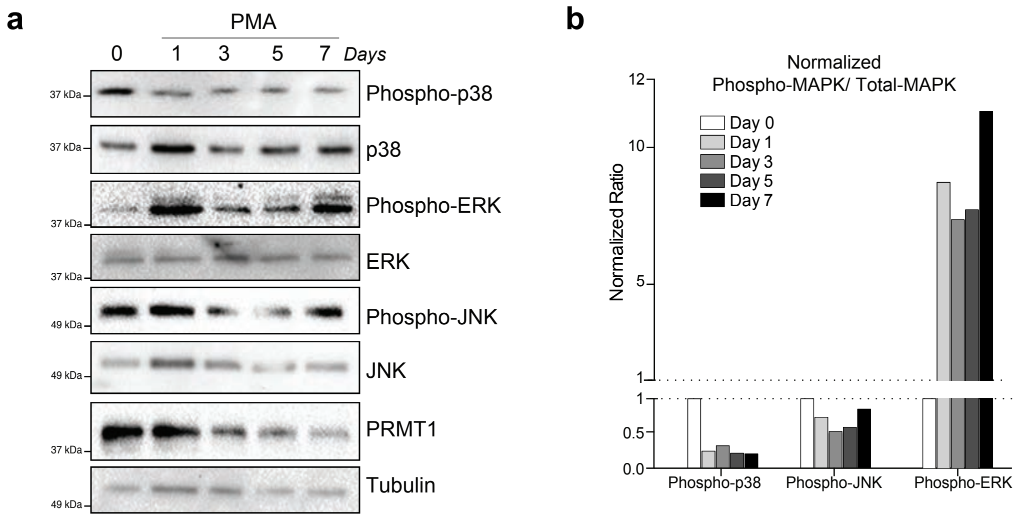

Su and Jiang et.al. Extended Data Fig. 3

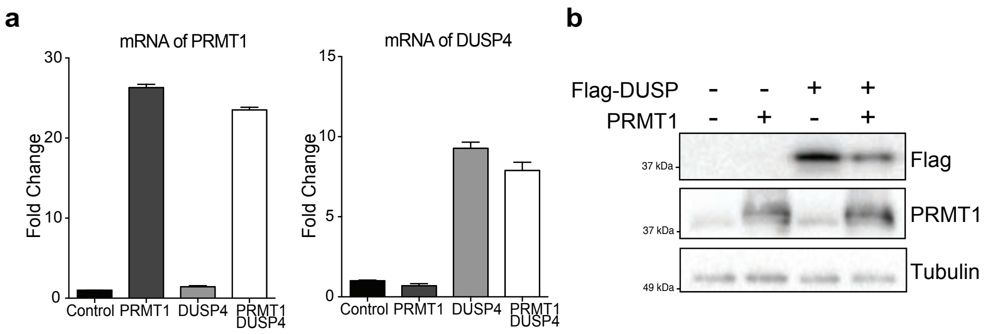

Su and Jiang et.al. Extended Data Fig. 4

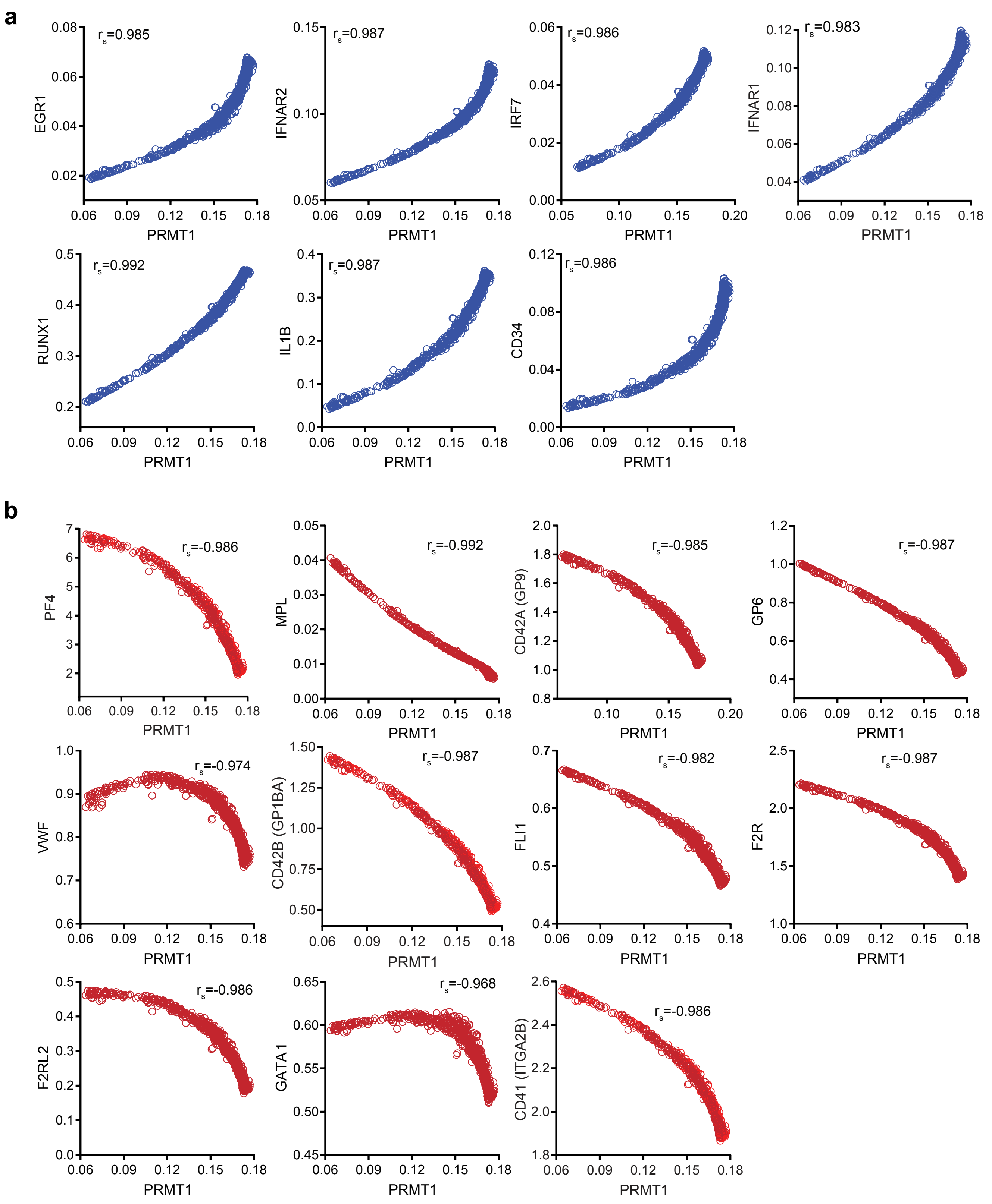

Su and Jiang et.al. Extended Data Fig. 5

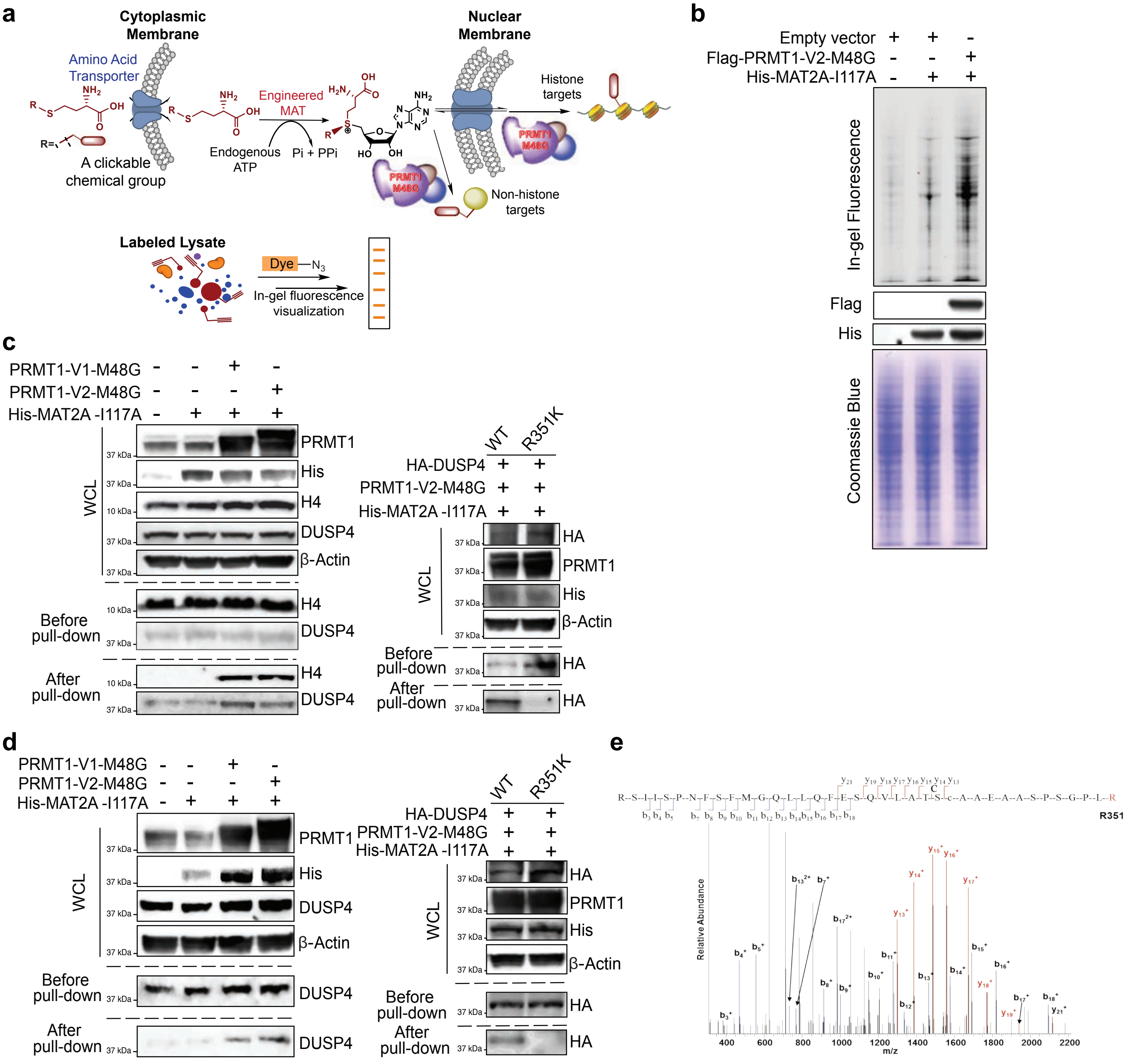

Su and Jiang et.al. Extended Data Fig. 6

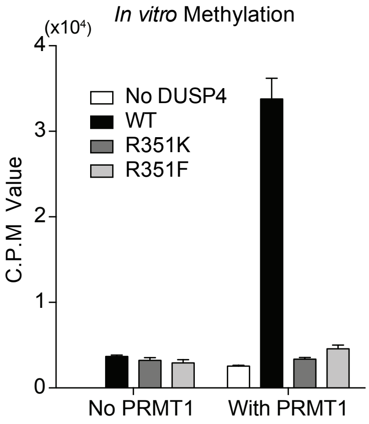

Su and Jiang et.al. Extended Data Fig. 7

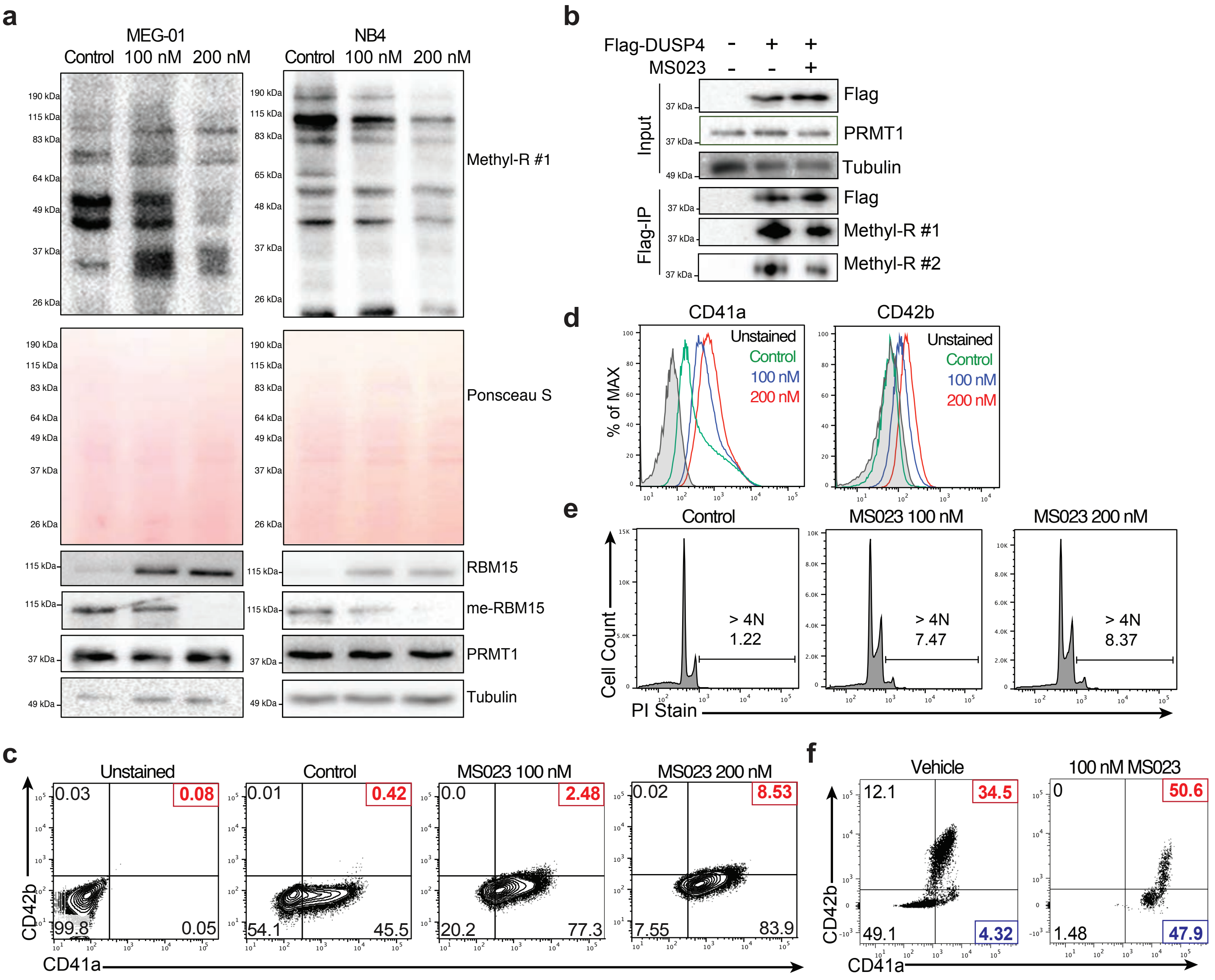

Su and Jiang et.al. Extended Data Fig. 8

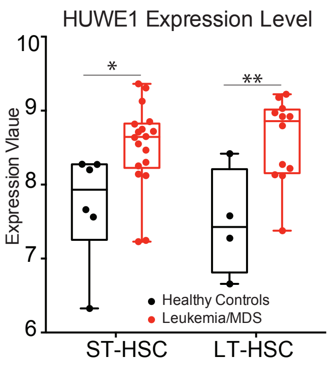
